# Multimodal Imaging Reveals Spatial Host-Pathogen Microenvironments in *Escherichia coli* Meningoencephalitis

**DOI:** 10.64898/2026.07.27.740408

**Authors:** Dominika Luptáková, Tereza Hřivnová Juříková, Oldřich Benada, Gabriela Lokočová, Helena Marešová, Hynek Mácha, Jiří Houšť, Kateřina Dvořáková Bendová, Miroslav Popper, Miloš Petřík, Andrea Palyzová, Lukáš Kučera, Evgeniya Biryukova, Jiří Novák, Vladimír Havlíček

**Affiliations:** Institute of Microbiology of the Czech Academy of Sciences, Vídeňská 1083, 142 00 Prague, Czech Republic; Institute of Molecular and Translational Medicine, Faculty of Medicine and Dentistry, Palacký University, Hněvotínská 5, 779 00 Olomouc, Czech Republic; Czech Advanced Technology and Research Institute, Palacký University, Šlechtitelů 241/27, 779 00 Olomouc, Czech Republic; Institute of Molecular and Translational Medicine, University Hospital, Hněvotínská 5, 779 00, Olomouc, Czech Republic; Institute of Molecular Genetics, Academy of Sciences of the Czech Republic, Vídeňská 1083, 142 00 Prague, Czech Republic; Department of Biochemistry, Faculty of Science, Charles University, Albertov 6, 128 00 Prague 2, Czech Republic; Department of Analytical Chemistry, Faculty of Science, Palacky University in Olomouc, 17. listopadu 1192/12, 779 00 Olomouc, Czech Republic

**Author notes:** **Corresponding authors:** Dominika Luptáková, and Vladimír Havlíček.

**Keywords:** Multimodal imaging, bacterial meningoencephalitis, *Escherichia coli*, siderophores, antimicrobial peptides, cerebrospinal fluid, spatial host-pathogen interplay

## Abstract

Experimental bacterial meningitis is often analyzed as microbial burden and inflammation, but the tissue-level organization of host-pathogen interplay remains poorly resolved. Here, we use multimodal imaging to map *Escherichia coli* meningoencephalitis as a spatially organized process across infected rat brain tissue and cerebrospinal fluid (CSF). A low-dose intracerebral inoculum expands within 24 h into an anatomically structured infection involving ventricular, meningeal, and perivascular interfaces. Scanning electron microscopy reveals biofilm-like multicellular *E. coli* aggregates embedded in extracellular material, consistent with neutrophil extracellular traps. MALDI mass spectrometry imaging detects aerobactin and salmochelin-derived metabolites, whereas intact enterobactin is not detected despite favorable analytical sensitivity. These bacterial iron-acquisition signals partially overlap with calprotectin proteoforms, defining ventricular and periventricular metal-conflict microenvironments. Spatial peptidomics identifies infection-enriched antimicrobial territories dominated by rat neutrophil peptides RatNP-2, RatNP-3, and RatNP-4. Endogenous proenkephalin-derived neuropeptides are reduced in basal ganglia regions. Finally, cerebrospinal fluid captures both bacterial siderophores and host antimicrobial peptides with lipocalin-2 protein, linking tissue-resolved host-pathogen chemistry to a proximal diagnostic fluid. Together, these data show that experimental *E. coli* meningoencephalitis is a spatially organized host-pathogen process. Bacterial communities, nutritional immunity, antimicrobial peptides, and neuropeptide networks occupy distinct but connected CNS niches, components of which are recoverable in CSF.

## Introduction

Bacterial meningitis remains a severe infectious disease in which mortality and neurological sequelae reflect both microbial invasion of the central nervous system (CNS) and the host inflammatory response [1]. Within this broad clinical entity, *Escherichia coli* is among the most common Gram-negative pathogens causing meningitis, particularly in neonates and infants [2], accounting for approximately 35% of clinical cases [3]. Meningitis-associated *E. coli* strains have evolved mechanisms that promote bloodstream survival, blood-brain barrier (BBB) crossing, and persistence in the CNS niche [4, 5]. Recent mechanistic work further supports BBB injury as a central event in neonatal *E. coli* meningitis [6]. Once bacteria enter the CNS, they encounter subarachnoid space, ventricular system, perivascular spaces, and further invade brain parenchyma via pial and glial interfaces [7]. These niches may influence whether *E. coli* remains mainly dispersed or forms multicellular, surface-associated communities. The tissue-level organization of bacterial cells after CNS entry remains insufficiently resolved.

Biofilms are multicellular bacterial communities embedded in extracellular polymeric matrices. In *E. coli*, biofilm matrix composition can include extracellular DNA (eDNA), extracellular curli amyloid fiber assembly, proteins, and polysaccharides, that serve as a structural and functional components [8], but the relative contribution of these components depends on strain, environment, and experimental conditions [9, 10]. Therefore, biofilm-like organization in infected tissue cannot be inferred from culture behavior alone and requires direct in situ visualization in the relevant tissue context.

The spatial organization of *E. coli* in the brain is likely coupled with bacterial iron acquisition and host metal-withholding responses [5, 11]. Pathogenic *E. coli* can deploy chemically distinct siderophore systems, including enterobactin, salmochelins and aerobactin [12]. These siderophores are not functionally interchangeable: salmochelin is a glucosylated enterobactin derivative, whereas aerobactin represents a chemically distinct iron-acquisition route [13]. Siderophore profiles might also be remodeled by degradation or processing, producing fragments of original molecules that retain low-affinity Fe-binding properties [14]. Bacteria have to cope with the nutritional immunity, in which host-specific proteins restrict their access to transition metals [11]. Key components of this response include lipocalin-2 (Lcn-2), targeting selected catecholate siderophores, e.g. enterobactin, but unrecognizing salmochelin, and neutrophil-derived calprotectin as an S100A8/S100A9 complex that sequesters nutrient metal ions in the inflammatory environments [15].

Neutrophil recruitment can also generate antimicrobial peptide-rich CNS microenvironments. Defensins are cationic host-defense peptides produced by neutrophils and epithelial cells, functioning in innate immunity beyond direct microbial killing [16]. Host-defense peptides have also been detected in the brain and spinal cord, where they may contribute to immunosurveillance and modulate host responses during infection and neuroinflammatory disease [17]. In rats, neutrophil peptides RatNP1-4 provide a molecular readout of α-defensin-based antimicrobial activity [18]. In parallel, CNS infection may perturb endogenous neuropeptide systems, including proenkephalin-derived peptides that are regionally organized in basal ganglia structures. Enkephalins are endogenous opioid peptides with established roles in CNS signaling, pain modulation, emotional regulation and neuroprotective or immunomodulatory contexts, but infection-associated spatial changes in proenkephalin-derived peptides remain poorly defined [19]. Thus, experimental *E. coli* meningoencephalitis provides an opportunity to examine bacterial metabolites and host inflammatory proteins, and the spatial relationship between antimicrobial peptides and endogenous neuropeptide networks.

Mass spectrometry imaging (MSI) is a central modality for spatially resolved multi-omics molecular mapping, enabling tissue-scale integration of metabolites, lipids, peptides and proteins [20] directly in tissue sections while preserving histological context [21]. Spatial multi-omics approaches illustrate how mass spectrometry imaging can be integrated with other imaging modalities to relate molecular heterogeneity to tissue architecture [22]. By combining optical microscopy, scanning electron microscopy (SEM), matrix-assisted laser desorption/ionization (MALDI) MSI-based metabolomics, peptidomics, and proteomics, with ELISA analysis, and liquid chromatography-mass spectrometry (LC-MS) of cerebrospinal fluid (CSF), for the first time, we connect tissue-spatially resolved host-pathogen microenvironments with fluid-accessible molecular signatures of CNS infection.

## Results

### Low-dose inoculum *E. coli* infection establishes anatomically structured meningoencephalitis

To establish an infection state suitable for spatial host-pathogen analysis, we compared two intracerebral *E. coli* inocula, 5˟10² colony forming unit (CFU) and 5˟10³ CFU, referred to low-dose and higher-dose inoculum model, respectively, with saline-injected controls. We analyzed brains 24 h after inoculation at +2.28 and -1.08 mm relative to bregma (Figure 1A and S1). The presence of bacteria in brain tissue was confirmed by Gram staining (Figure S1A), quantitative PCR and culture. After 24 h, *E. coli* burden in a low-inoculum model reached up to 1.4˟10⁵ genome equivalents/mg tissue by quantitative polymerase chain reaction (qPCR) and 6.1˟10³ CFU/mg tissue by culture (Figure 1B). Histopathological examination of hematoxylin and eosin (HaE)-stained brain sections demonstrated a reproducible inflammatory phenotype. In the low-dose inoculum model, infected brains exhibited more pronounced leptomeningeal (*P* < 0.0001) and perivascular (*P* = 0.0003) inflammation, neutrophil-rich ventriculitis (*P* = 0002), and reactive gliosis (*P* = 0.0016) compared to controls, suggesting a slower infection dynamic with an organized immune response centered in the meninges and around blood vessels (Figure 1C,D and S1C). In contrast, the higher-dose inoculum model showed predominant parenchymal involvement, vascular injury, and ventriculitis, consistent with rapid bacterial dissemination and acute tissue destruction (Figure S1B,C). These findings indicate that the inflammatory profile and localization within the CNS depend on infection intensity and on the tempo of host-pathogen interaction. Because higher-inoculum phenotype approached overt tissue destruction, whereas the low-dose inoculum model preserved a spatially interpretable inflammatory architecture, subsequent SEM and MALDI MSI analyses were performed using the latter model.

**Figure 1.**
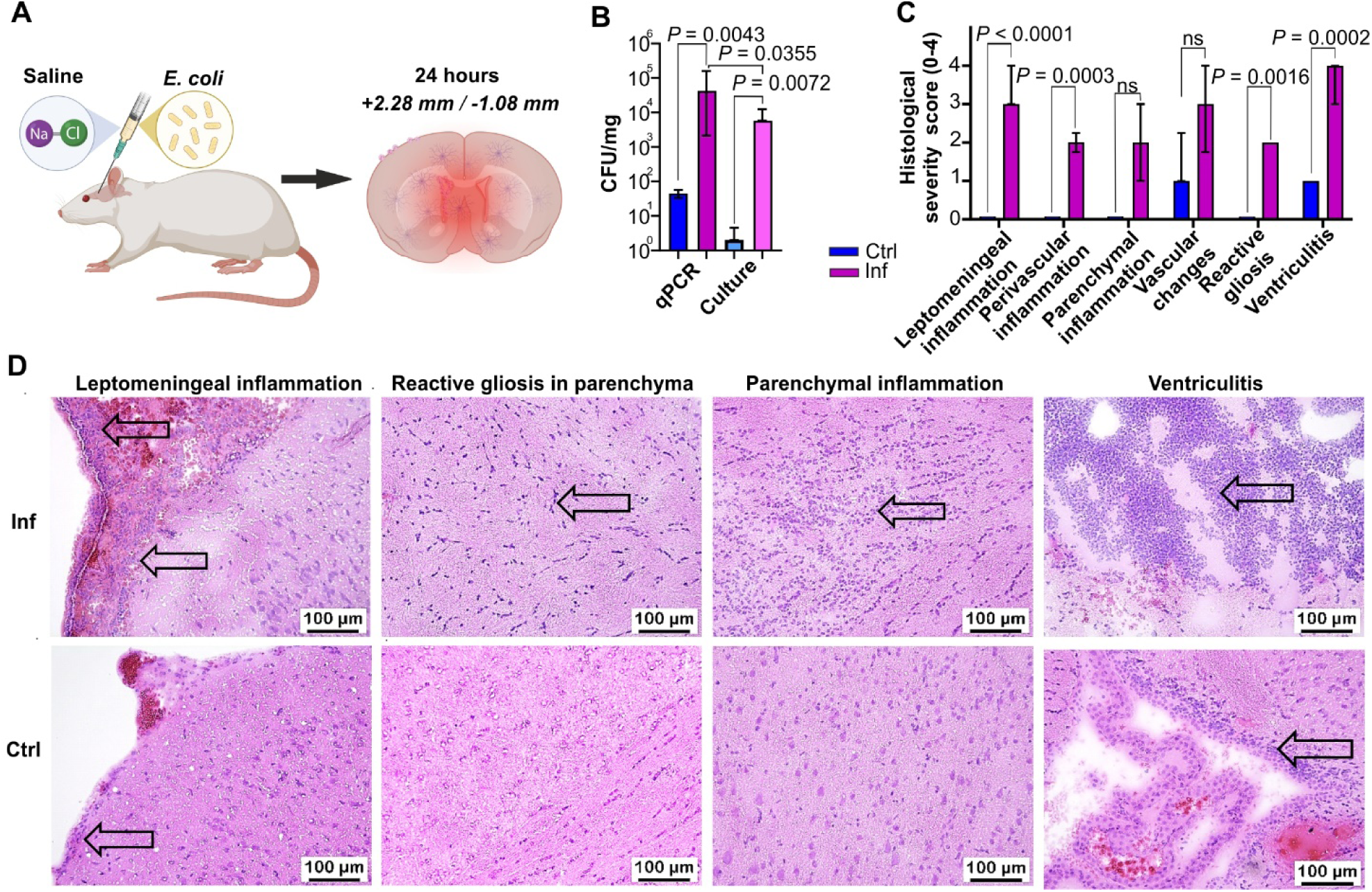
Experimental model of low-dose inoculum *E. coli* showing anatomically structured meningoencephalitis. (A) Experimental design. Rats received intracerebral saline or *E. coli* inoculation and brains were analyzed 24 h later at the tissue level of +2.28 mm and -1.08 mm relative to bregma. (B) *E. coli* burden in brain tissue from control (*n* = 4) and infected (*n* = 5) animals measured by qPCR and culture in the low-dose inoculum (5˟10² CFU) model. qPCR and culture data are expressed as genome equivalents per mg tissue and CFU per mg tissue, respectively. (C) Histopathological scoring of CNS pathology in control and infected brains using a 0-4 severity scale evaluated at both tissue levels. Differences in bacterial burden and histopathological scores between control and infected groups were assessed using the Kruskal–Wallis test followed by an uncorrected Dunn’s multiple-comparisons test. Data are shown as median ± interquartile range. Exact *P* values are indicated in the graphs. (D) Representative hematoxylin and eosin-stained sections showing neuropathology, indicated by arrows, in infected brains, with corresponding control regions shown below. Scale bars, 100 μm. qPCR, quantitative polymerase chain reaction; CFU, colony forming unit; Inf, infection; Ctrl, control; ns, not significant.

### Scanning electron microscopy reveals biofilm-like *E. coli* communities at CNS tissue interfaces

In the established low-dose inoculum model, we investigated the morphological organization of *E. coli* cells and immune response within infected brain tissue. High-resolution SEM of post-fixed fresh-frozen sections showed bacterial clusters in tissue areas exposed to ventricular and inner brain interfaces (Figure 2A,B). Bacteria were detected as individual rod-shaped cells and as surface-associated multicellular aggregates within cavities and along tissue surfaces. Their distribution along with ventricular and gyri structures was consistent with access to CSF-facing anatomical compartments. At higher magnification, the aggregates showed a biofilm-like architecture, with tightly apposed bacterial cells embedded in abundant extracellular material (Figure 2C). Most bacterial cells retained the morphology of Gram-negative bacilli, whereas subsets showed deformation or altered surface contours, indicating morphological heterogeneity within the tissue-associated bacterial population (Figure 2D). A prominent feature of infected regions was an extracellular fibrillar scaffold, dense mesh-like structures, surrounding and interconnecting bacterial aggregates (Figure 3A). The thick, densely packed layer of bacterial aggregates were interconnected by extracellular tubular fibers approximately of 200 nm in thickness, pointing to the neutrophil membrane tubulovesicular extension, and some were trapped on thin filaments about 20 nm thick, comparable to the thickness of neutrophil traps (Figure 3B,C). SEM further revealed bacterial surface-associated features, including fine pili-like fibers and thicker tubular membrane protrusions. The two types of filamentous structures occurred in the infected tissue, including thin filaments suggesting the neutrophils traps, and the thicker fibrin network (Figure 3D). The spatial association of bacterial biofilm-like aggregates with extracellular neutrophil tubular extensions corresponded to the presence of bacteria and neutrophils cells in the same brain region as shown by histology (Figure 1D and S1A).

**Figure 2.**
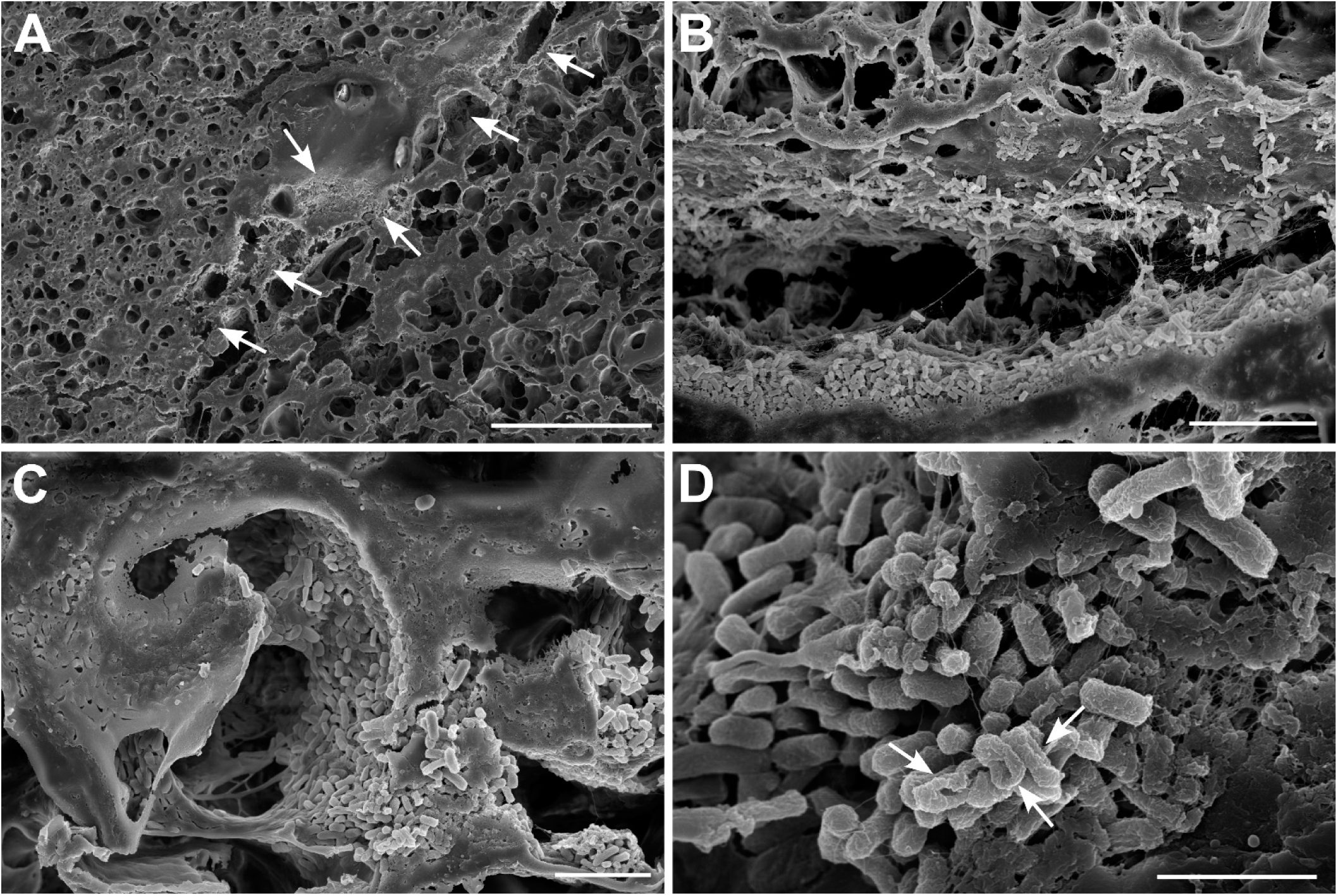
High-resolution SEM revealing biofilm-like *E. coli* communities in infected brain tissue. (A) Low-magnification SEM overview of infected brain tissue section showing bacterial accumulation along tissue cavities and inner brain surfaces. Arrows indicate regions containing bacterial cells or aggregates. (B) Higher-magnification view showing individual bacilli and bacterial clusters. (C) Bacterial biofilm on the inner brain surfaces. (D) High-magnification view of a biofilm-like bacterial community. Most cells retain rod-shaped Gram-negative shape, whereas others show altered morphology or deformation, as indicated by arrows. Scale bars: A = 50 μm, B = 10 μm, C = 5 μm, D = 2 μm.

**Figure 3.**
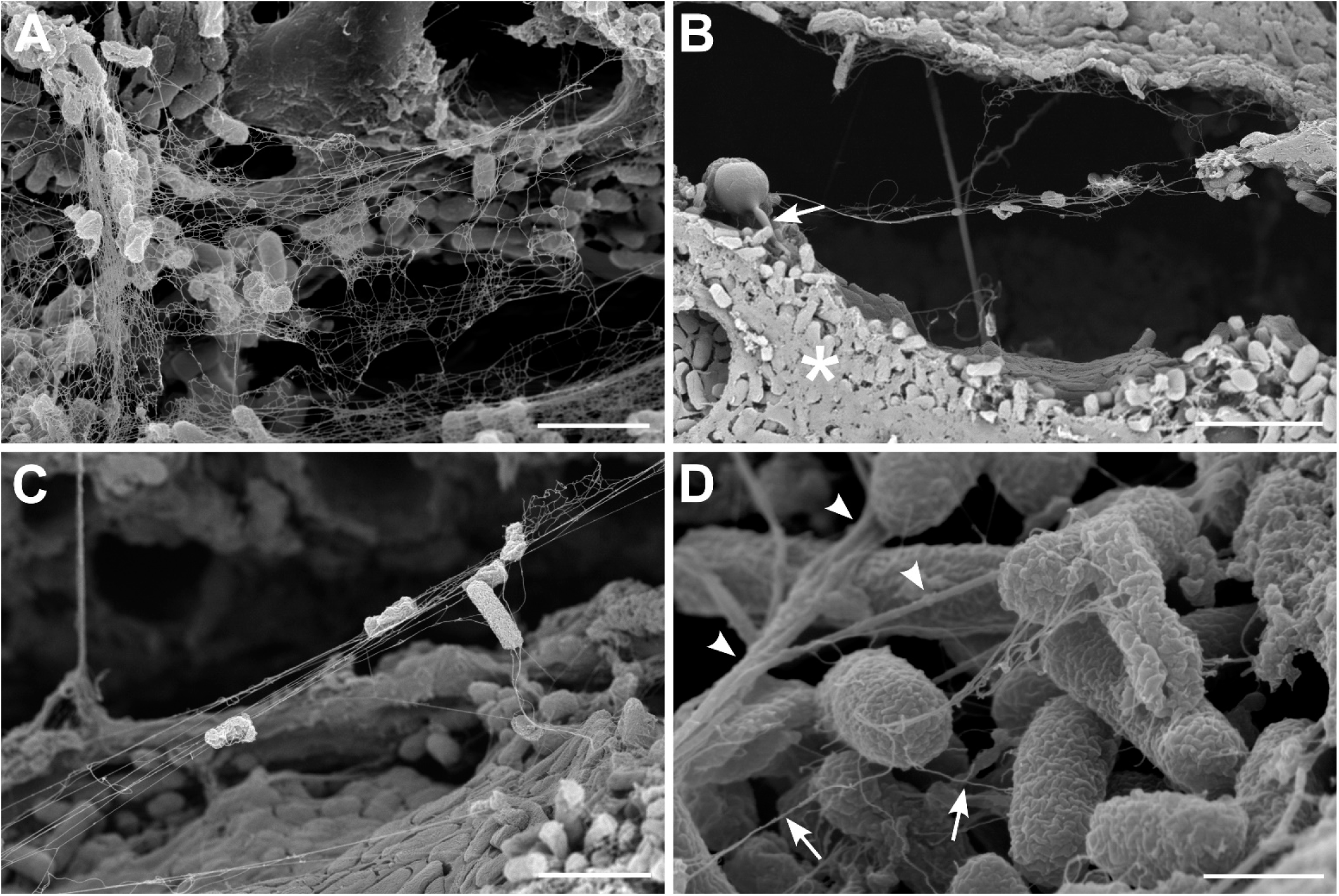
High-resolution SEM showing filamentous structures exclusively present in infected brain tissue. (A) Dense mesh-like structures heavily covering the bacterial cells. (B) A cross section of the thick, densely packed bacterial biofilm layer marked by an asterisk. The arrow points to a tubular structure resembling a neutrophil membrane tubulovesicular extension, with trapped bacteria. (C) Higher magnification showing trapped bacteria. (D) High magnification of the top layer of the bacterial biofilm showing well-preserved *E. coli* outer membrane surface. The panel shows the two types of filamentous structures occurring in the sample. The thin filaments plausibly represent neutrophil traps (arrows), while the thicker ones represent the fibrin network (arrowheads). Scale bars: A = 2 μm, B = 4 μm, C = 2 μm, D = 0.5 μm.

### Bacterial siderophores and host calprotectin define partially overlapping metal-conflict niches

Using MALDI MSI we examined spatial organization of bacterial and host metal sequestration molecular systems in the CNS tissue interfaces. We mapped *E. coli* siderophores and the neutrophil-associated calprotectin subunits S100A8 and S100A9 at +2.28 mm and -1.08 mm tissue level relative to bregma (Figure 4 and S3). The *E. coli* CCM 5172 strain produced aerobactin, enterobactin and salmochelin under iron-limited conditions in vitro (data not shown). In infected brain tissue, we detected aerobactin as the predominant siderophore signal, together with salmochelin-derived metabolites, mainly salmochelin SX and 2,3-dihydroxybenzoylserine (DHBS) (Figure 4A-C and Table S1). Enterobactin was not detected in infected sections, despite a lower tissue detection limit for enterobactin than for aerobactin in the MALDI quantitative MSI (qMSI) validation dataset (Figure S2 and Table S2). In parallel, we displayed calprotectin-related signals of S100A8, acetylated S100A9 (S100A9-ac) and acetylated/methylated S100A9 (S100A9-ac/m) proteoforms. Among them, S100A9-ac/m generally showed the highest regional signal, followed by S100A9-ac and S100A8 (Figure 4B-D and S3, and Table S3). As an additional host nutritional-immunity marker, Lcn-2 was exclusively detected in infected brain tissue (Figure. S4).

**Figure 4.**
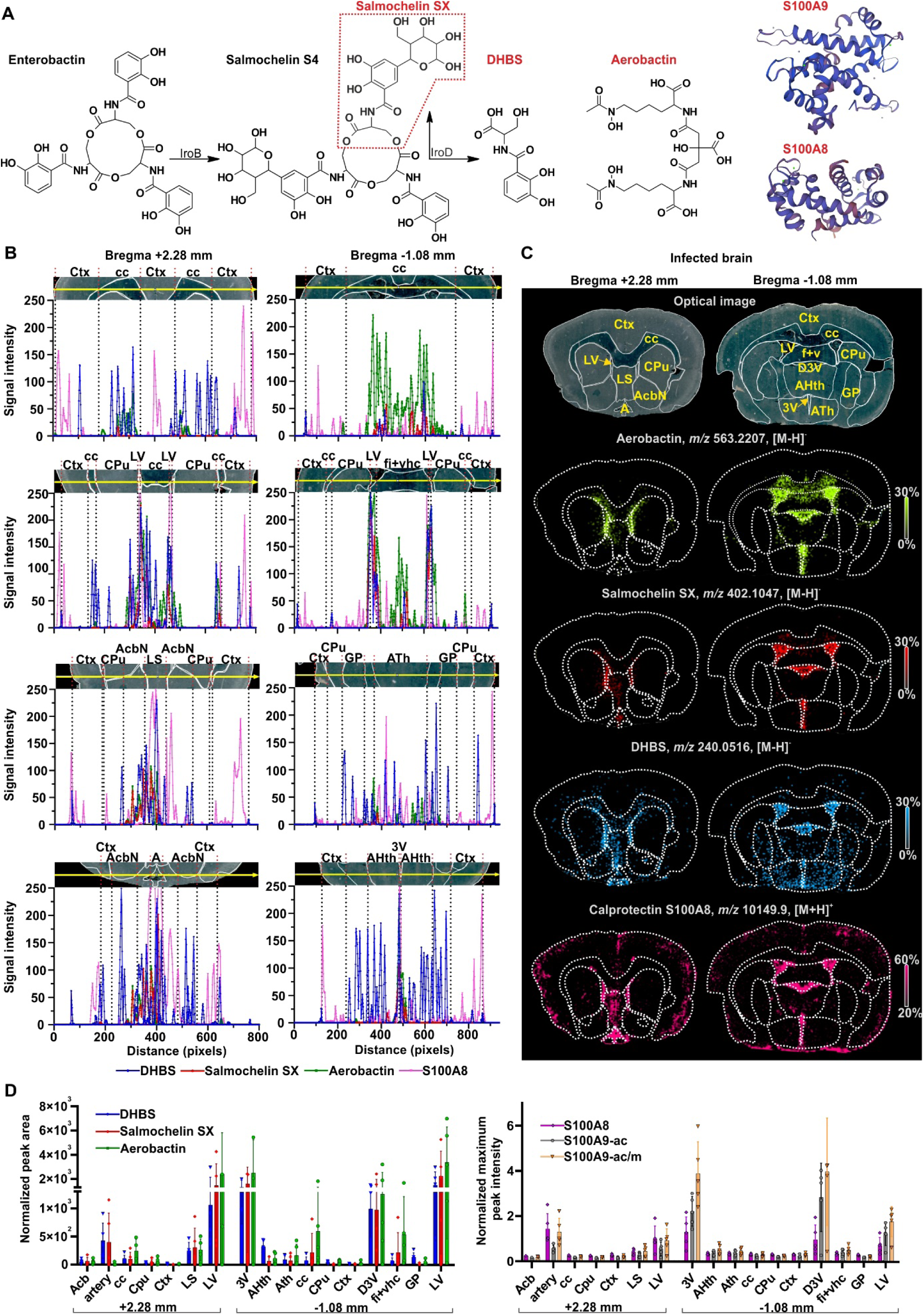
Bacterial siderophores and calprotectin defining metal-withholding niches in *E. coli*-infected brain tissue at +2.28 mm and −1.08 mm level relative to bregma. (A) Schematic representation of *E. coli* siderophores, including their synthetic relationships, and the host calprotectin subunits S100A9 and S100A8. Detected compounds by MALDI MSI are shown in red. (B) Line-scan profiles showing regional signal distributions of 2,3-dihydroxybenzoylserine (DHBS), salmochelin SX, aerobactin, and S100A8 across selected anatomical structures in infected brain sections. Dashed lines indicate region boundaries. Optical images above each plot show the anatomical regions used for line-scan analysis. The yellow line in panel indicates the line-scan trajectory. (C) Representative MALDI MSI ion maps of aerobactin, salmochelin SX, DHBS and S100A8 in infected brain sections. Siderophore signals were root-mean-square normalized; S100A8 signal was total-ion-current normalized. Ion images are displayed on linear intensity color-coded scales. (D) Quantification of normalized peak areas of siderophores (left panel) and normalized maximum peak intensities of calprotectin proteoforms (right panel) across brain regions. Data are presented as mean ± standard deviation (*n* = 5). A, cerebral artery; LS, lateral septum; Ctx, cortex; CPu, caudate putamen; AcbN, accumbens nucleus; cc, corpus callosum; LV, lateral ventricle; D3V, dorsal third ventricle; 3V, third ventricle; ATh, anterior thalamus; AHth, anterior hypothalamus; fi+vhc (f+v), fimbria of hippocampus and ventral hippocampal commissure; GP, globus pallidus; ac, acetylation; m, methylation.

Correlation analysis revealed positive associations between bacterial siderophores and calprotectin proteoforms (Table S4). At -1.08 mm brain level, most consistently DHBS positively correlated with calprotectin-related signals S100A8 (ρ = 0.52, *P* = 0.0039, FDR q = 0.0171), S100A9-ac (ρ = 0.54, *P* = 0.0024, FDR q = 0.0171), and S100A9-ac/m (ρ = 0.53, *P* = 0.0032, FDR q = 0.0171). At +2.28 mm, the strongest associations involved DHBS with S100A9-ac/m (ρ = 0.78, *P* = 0.0048, FDR q = 0.0149) and S100A8 (ρ = 0.70, *P* = 0.0003, FDR q = 0.0033). Partial concordance became clearer when the regional host-siderophore balance was examined. By using line analysis of ion maps we determined that siderophores and calprotectin signals were enriched around the ventricular axis and adjacent tissue interfaces, however, their local maxima differed (Figure 4B,D). Host-dominant regions included the cerebral artery at +2.28 mm, the dorsal third ventricle at -1.08 mm, and brain cortex at both levels. Lateral ventricular regions, corpus callosum, and fimbria of hippocampus and ventral hippocampal commissure region showed relatively stronger siderophore enrichment (Figure. 4B-D, Tables S5).

### Unsupervised spatial peptidomics identifies infection-enriched antimicrobial peptide territories

It is increasingly clear that the nervous and immune systems use intersystem communication on molecular level via peptides [23]. MALDI MSI spatial peptidomics detected 111 tissue-associated molecular features, of which 20 were assigned to endogenous peptides, including proenkephalin-derived peptides, cerebellin-1 precursor fragments, ProSAAS-derived peptides, thymosin β4, thymosin β10 and rat neutrophil antimicrobial peptides (RatNPs) (Table S6). Several assigned peptides carried post-translational modifications, including acetylation, phosphorylation or disulfide bonds. Uniform Manifold Approximation and Projection (UMAP) followed by bisecting k-means segmentation across control and infected sections from both bregma levels resolved six molecularly distinct spatial clusters (Figure 5A,B). Control tissue was dominated by clusters enriched in endogenous lipid- and peptide-associated features, whereas infected sections contained territories with prominent signals in the *m/z* range corresponding to peptides annotated as RatNP-2-4 (Figure 5B,C). Cluster-resolved quantification showed the RatNPs were concentrated around ventricular, limbic, and meningeal/peripheral brain regions (Figure 5D). These territories partially overlapped with infection-enriched regions (Figure 4), showing the decreasing distribution trend towards the adjacent regions. Control-enriched clusters contributed little to the total RatNP signal (Figure 5D). Targeted quantification within infected tissues confirmed strong RatNP signals at both anatomical levels, with the lowest abundance of RatNP-2 compared to a high level of RatNP-3 (*P* = 0.0047 at +2.28 mm and -1.08 mm level) and RatNP-4 that displayed a comparable abundance (Figure 5E).

**Figure 5.**
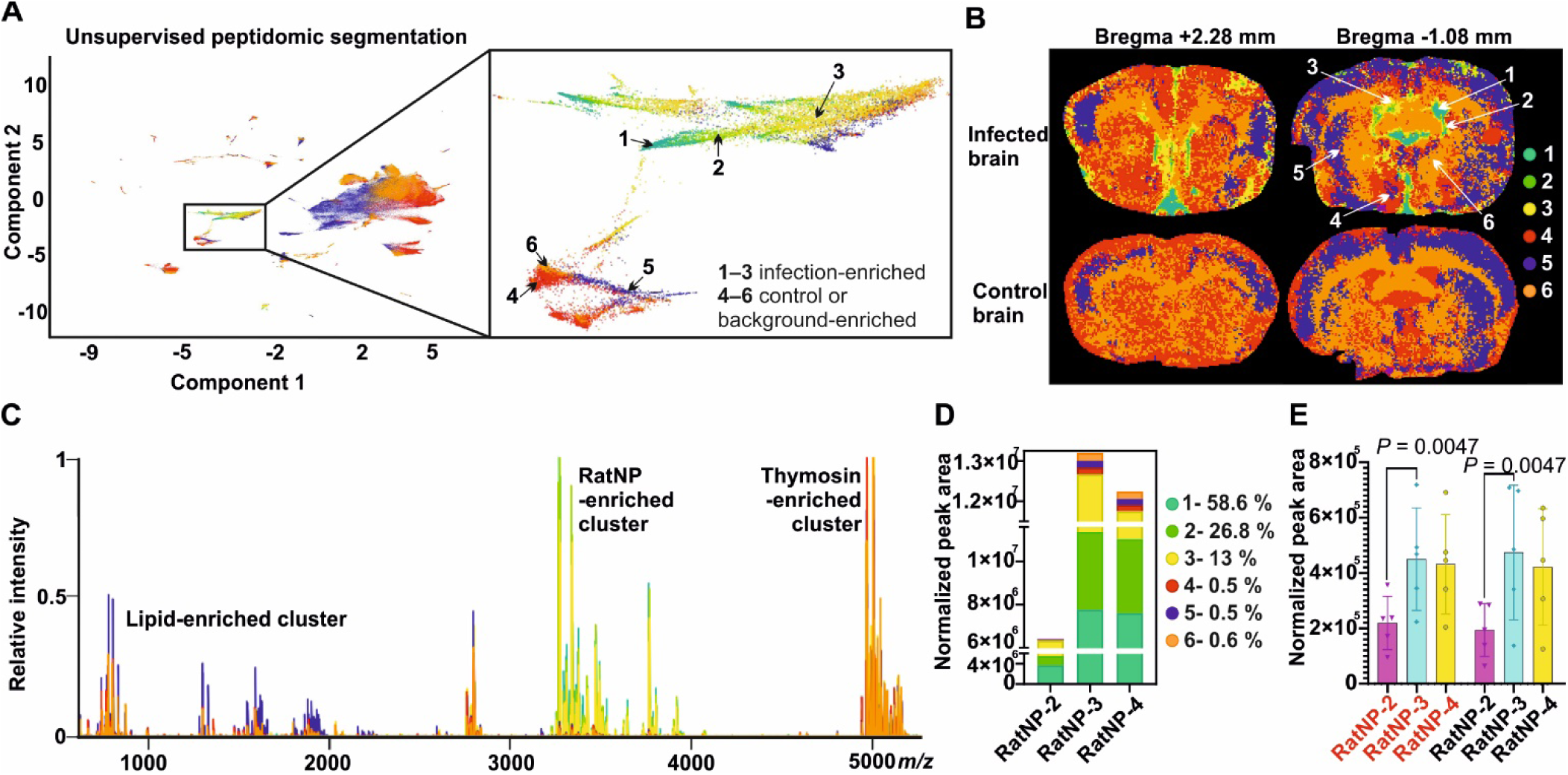
Unsupervised spatial peptidomics revealing infection-specific molecular zones. (A) UMAP projection of MALDI MSI peptidomic data considering 111 tissue-associated molecular features present in control (*n* = 4) and infected (*n* = 5) brain sections at +2.28 mm and -1.08 mm relative to bregma. UMAP was followed by bisecting k-means segmentation, resolving six molecularly distinct clusters displayed in zoom. (B) Spatial projection of the six peptidomic color-coded clusters onto representative infected and control brain sections at both anatomical levels. (C) Representative spectrum of overlapping six cluster-related mean mass spectra. Infection-enriched clusters 1-3 showed prominent signals in the *m/z* range of rat neutrophil peptides. Clusters 4-6 were dominated by lipid-associated or endogenous peptide features, including thymosin-related signals. (D) Percentual contribution of the total RatNP-2, RatNP-3 and RatNP-4 signal to individual clusters. (E) Relative quantification of RatNP-2, RatNP-3 and RatNP-4 abundance in infected tissue sections (*n* = 5) at +2.28 mm (red titles on x-axis) and -1.08 mm level relative to bregma. Data are expressed as mean ± standard deviation. Statistical testing was performed separately for each anatomical level using Friedman’s test followed by Dunn’s multiple comparisons test. Exact *P* values are indicated.

### Rat neutrophil peptides define ventricular and meningeal antimicrobial zone gradient in the infected brain

Supervised analysis of peptide distributions defined significant differences in the abundance of individual peptides between control and infected tissues (Figure 6A, Table S6). RatNPs increased in infected brains by 181- to 226-fold at +2.28 mm and by 860- to 2303-fold at -1.08 mm relative to bregma, indicating a robust neutrophil-derived antimicrobial peptide response during *E. coli* meningoencephalitis. Spatial analysis showed that RatNPs were not uniformly distributed across infected brain tissue. We suggest that the RatNP signal exhibit directional propagation, extending from the meninges inward into the brain parenchyma and from central brain regions toward the periphery (Figure 6B-D). Line-scan image analysis revealed that RatNP-2-4 followed highly similar spatial distributions. RatNP signals peaked at brain cortex, ventricular and interface-associated compartments localized around brain midline, and decreased in adjacent parenchymal regions, e.g. caudate putamen and accumbens nucleus (Figure 6C,D). Ventricles, including the lateral ventricles, the third ventricle, and the dorsal third ventricle, as well as the basal arterial region, showed the strongest RatNP enrichment. RatNP-3 was the dominant peptide species reaching the concentration up to 30.6 µg/g of tissue in the highest-abundance regions. Intermediate RatNP-3 levels were determined in the lateral septum (9.9 µg/g), anterior thalamus (5.3 µg/g), and fimbria of hippocampus and ventral hippocampal commissure (6.8 µg/g), whereas other regions contained RatNPs at low abundance or near the limit of detection (Figure 6E and Table S7). Additionally, at both anatomical levels, RatNPs significantly correlated with S100A8 and S100A9 proteoforms, particularly RatNP-2 (at 2.28 mm: ρ = 0.93, *P* = 0.0007, FDR q = 0.0043 for S100A8; ρ = 0.93, *P* = 0.0002, FDR q = 0.0033 for S100A9-ac; at -1.08 mm: ρ = 0.95, *P* = 0.0001, FDR q = 0.0022 for S100A9-ac) (Table S4). In contrast, we determined a heterogenous correlation between RatNPs and siderophores. RatNPs correlated most consistently with DHBS, significantly with RatNP-2 (ρ = 0.81, *P* = 0.0044, FDR q = 0.0149 at +2.28 mm; ρ = 0.53, *P* = 0.0047, FDR q = 0.0171 at -1.08 mm), whereas correlations with aerobactin and salmochelin SX were weaker and varied between anatomical levels (Table S4).

**Figure 6.**
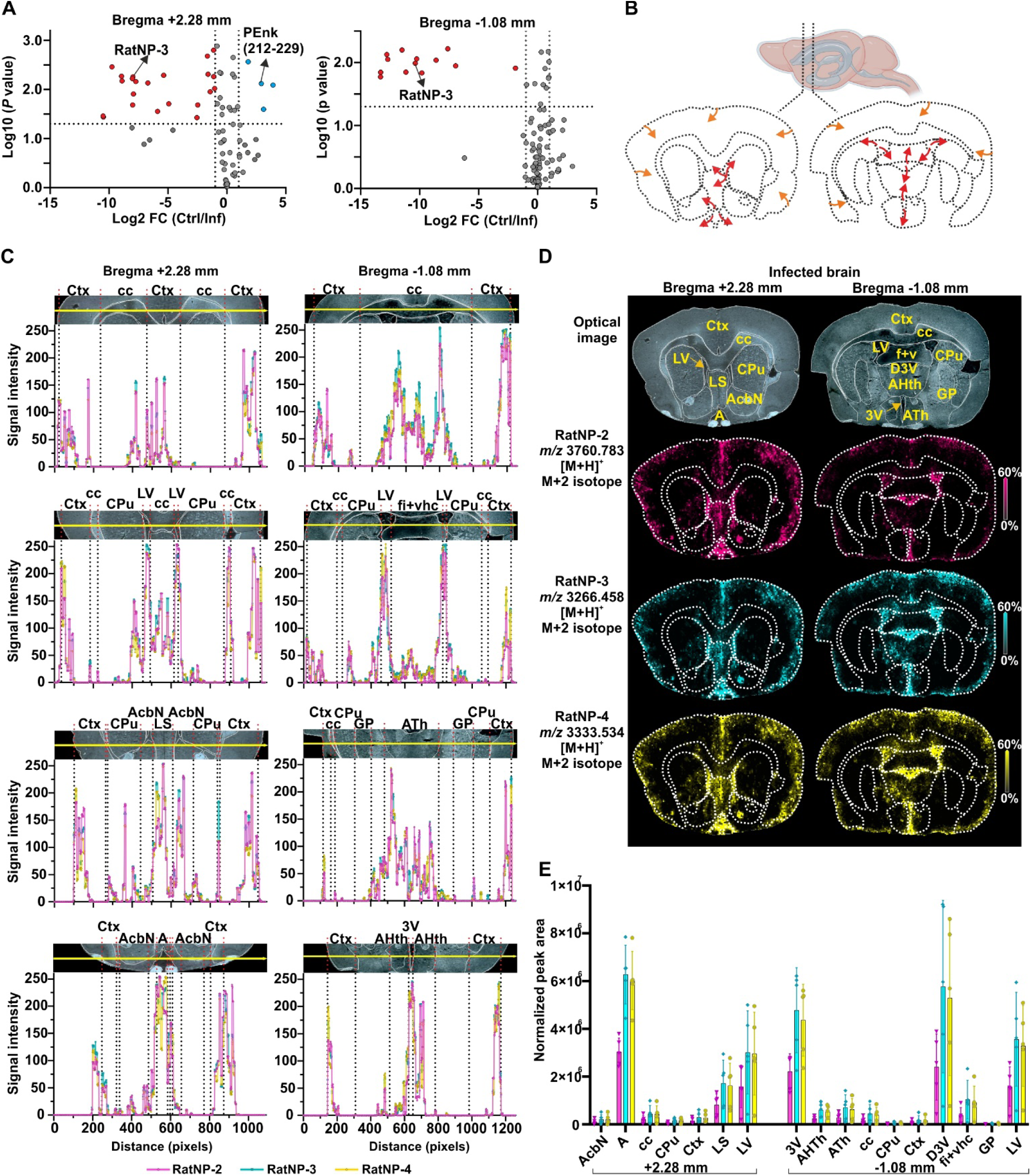
Spreading of rat neutrophil peptides in *E. coli*-infected brain tissue at +2.28 mm and - 1.08 mm relative to bregma. (A) Volcano plots showing differential abundance of 111 MALDI MSI-detected molecular features between control (Ctrl, *n* = 4) and infected (Inf, *n* = 5) brain sections. Significantly increased and decreased features in infected tissue are shown in red and blue, respectively. Dashed lines indicate *P* ≤ 0.05 and fold change (FC) > 2. (B) Schematic summary of the anatomical distribution of RatNP signals based on regional MALDI MSI analysis. Arrows indicate suggested diffusion of the RatNPs signal from meninges into the brain parenchyma (orange) and from ventricles into the surrounding regions (red). (C) Line-scan intensity profiles of RatNPs-2-4 across selected brain regions. Dashed lines indicate region boundaries. Optical images above each plot show the anatomical regions used for line-scan analysis. The yellow line in panel indicates the line-scan trajectory. Note identical spatial distribution of all RatNPs. (D) Representative optical images and MALDI MSI ion maps showing the spatial distributions of RatNP-2-4 in infected brain sections. Mouse hepcidin 25-normalized ion abundances are displayed on linear color-coded scales. (E) Region-resolved quantification of normalized RatNP-2-4 abundance across defined brain regions in infected animals (*n* = 5). Data are expressed as mean ± standard deviation. A, cerebral artery; LS, lateral septum; Ctx, cortex; CPu, caudate putamen; AcbN, accumbens nucleus; cc, corpus callosum; LV, lateral ventricle; D3V, dorsal third ventricle; 3V, third ventricle; ATh, anterior thalamus; AHth, anterior hypothalamus; fi+vhc (f+v), fimbria of hippocampus and ventral hippocampal commissure; GP, globus pallidus.

### *E. coli* meningoencephalitis reduces the basal ganglia proenkephalin peptide network

In MALDI MSI data, we annotated ten proenkephalin-derived peptides, assigned by accurate mass and by their spatial distributions, which matched their expected enrichment in basal ganglia structures, particularly the globus pallidus followed by caudate putamen and nucleus accumbens (Figure 7 and Table S6). Compared to controls, infected brains showed a coordinated reduction in PEnk-derived signals. Infection was associated with a 49% lower total PEnk signal relative to controls (*P* = 0.0293; Figure 7A,B). The most significant decreases were observed for PEnk (143–185) (*P* = 0.0148) and PEnk (219–229) (*P* = 0.0051), corresponding to a 47% signal decrease across both analyzed tissue levels (Figure 7C). The reduction was also evident within the globus pallidus (Figure 7B,C). This points to a broad suppression of the proenkephalin system, extending beyond individual peptide alterations.

**Figure 7.**
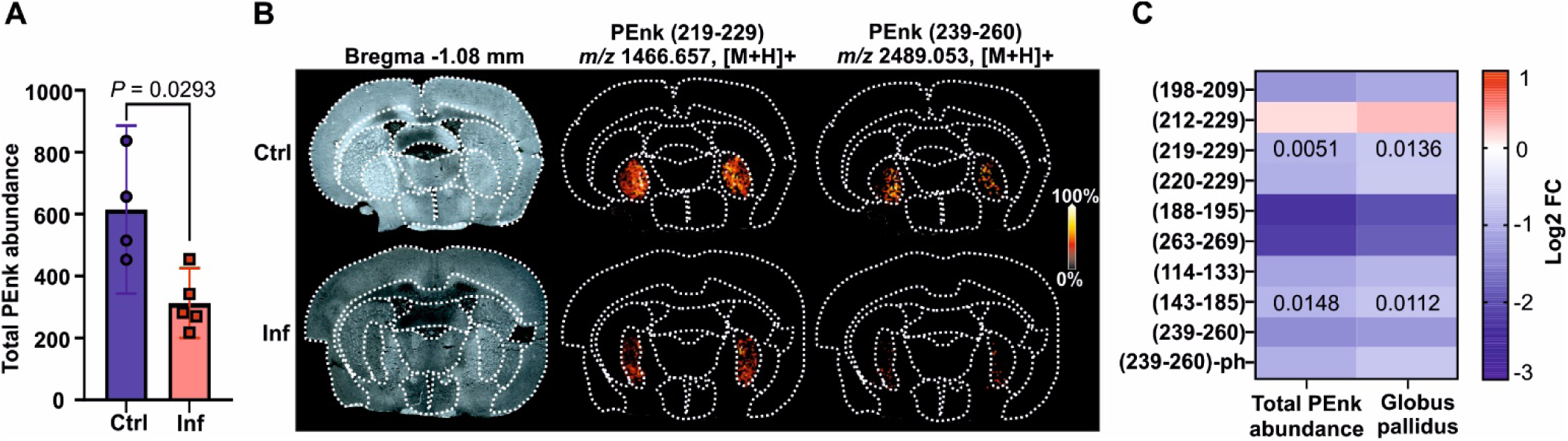
Altered proenkephalin (PEnk) peptide level induced by *E. coli* meningoencephalitis. (A) Total PEnk abundance in control (Ctrl, *n* = 4) and infected (Inf, *n* = 5) brain tissue sections, determined as the mean of summed PEnk signals detected at both +2.28 mm and −1.08 mm levels relative to bregma for each animal. Bars show mean ± standard deviation. Statistical significance was assessed using Welch’s t test. (B) Representative optical images and MALDI MSI ion maps showing PEnk (219-229) at *m/z* 1466.657 [M+H]^+^ and PEnk (239-260) at *m/z* 2489.053 [M+H]^+^ in control and infected brain sections at -1.08 mm relative to bregma. Ion images are displayed as root-mean-square-normalized ion abundance on a linear intensity color-coded scale. (C) Heatmap showing differential abundance of detected PEnk-derived peptides in infected versus control brain tissue. Colors indicate log2 fold change (FC) (Inf vs. Ctrl). Numbers indicate exact *P* values for peptides reaching statistical significance in the total abundance-level analysis or within the globus pallidus region. Statistical comparisons were performed using unpaired t test with Welch correction, followed by Bonferroni-Dunn’s multiple comparisons test.

### CSF captures host antimicrobial peptides with lipocalin-2 and bacterial siderophores during E. coli meningoencephalitis

As a translational step towards clinical diagnostics, we analyzed CSF from control and infected animals for determined host-pathogen molecular signatures using MALDI-MS and ELISA (Figure 8). Antimicrobial peptides RatNP-2, RatNP-3, and RatNP-4, and protein Lcn-2 were detected in CSF from infected animals, whereas they remained undetectable in control CSF samples (Figure 8A, B). RatNP-3 and RatNP-4 showed the highest relative abundances, while RatNP-2 was detected at lower levels, broadly consistent with the antimicrobial peptide pattern observed in tissue. LC-MS analysis further detected multiple *E. coli* siderophores exclusively in infected CSF. Aerobactin was the most abundant siderophore signal, accompanied by salmochelin-derived metabolites, including salmochelin S1, S2, S5 and salmochelin SX (Figure 8C). Note, enterobactin was not detected in CSF, similarly to tissue analysis. In control samples, siderophore signals were below limit of detection.

**Figure 8.**
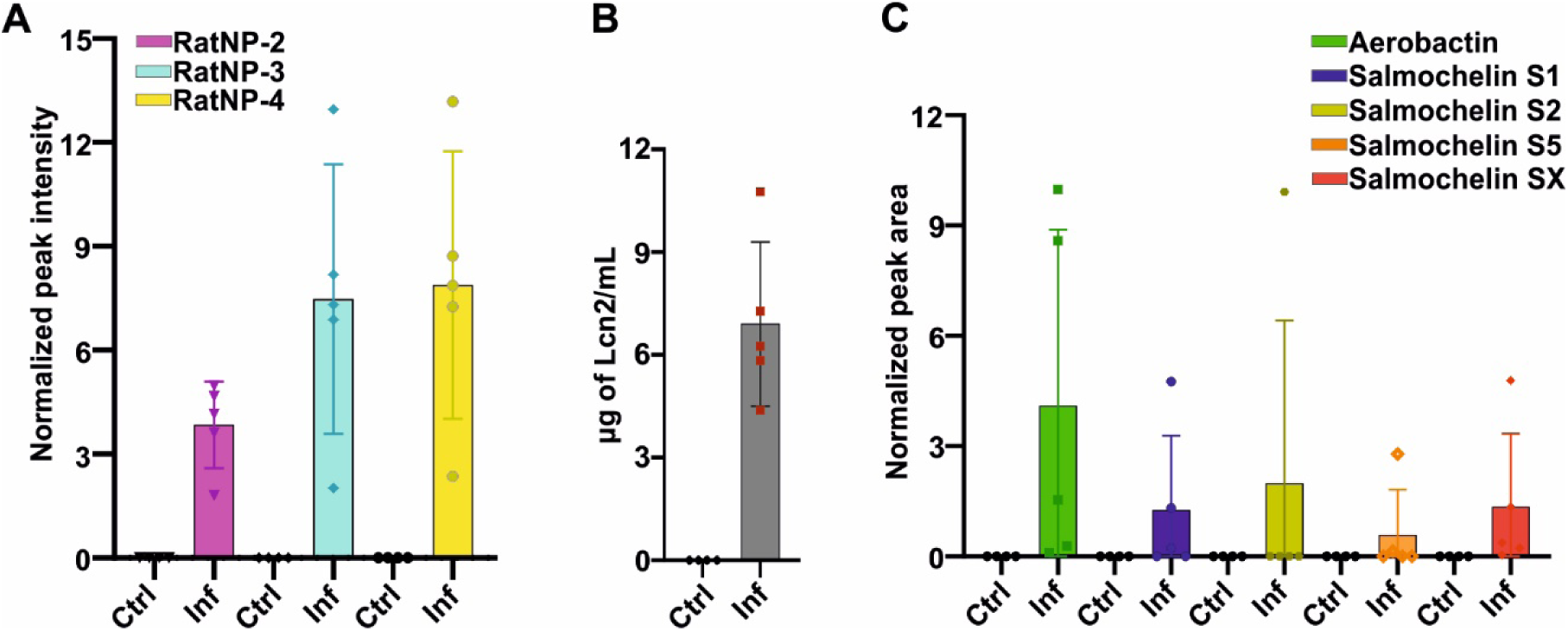
Analyses of cerebrospinal fluid for the host-pathogen-derived molecular markers. (A) Relative quantification of rat neutrophil peptides RatNP-2, RatNP-3 and RatNP-4 by MALDI MS. (B) ELISA-based quantitation of lipocalin-2 (Lcn-2). (C) Relative quantification of *E. coli* siderophores by LC-MS. Samples analyzed originated from control (*n* = 4) and infected (*n* = 5) animals. Data are shown as mean ± standard deviation.

## Discussion

Bacterial meningitis and meningoencephalitis are understood as invasion of meningeal and perivascular space by bacteria crossing BBB and blood-CSF barriers and causing an inflammatory injury in CSF, meninges, and brain parenchymal compartments [24]. Current comprehension has been shaped largely by bacterial burden and inflammatory mediators [25], which, being averaged across tissue, cannot resolve how the participating host- and pathogen-specific molecules are arranged. By combining SEM with MALDI MSI-based spatial metabolomics, peptidomics and proteomics on serial sections of the same brains, we define *E. coli* meningoencephalitis as a spatially organized morphological and molecular host-pathogen process. A few studies have systematically mapped how bacteria are organized once inside brain tissue. Henke et al. [26] showed by transmission electron microscopy that *Listeria monocytogenes* are located within neuronal axons, which is accompanied by axon swelling and tightening of myelin sheath. Other studies using immunofluorescence showed *Streptococcus pneumoniae* adhere preferentially to the subarachnoid vessels and reach the more internal cerebral areas including the cerebral cortex, septum, choroid plexus [27], and hippocampus [28]. Using the float-fixation of cryosections [29], we revealed that E. coli, both as individual bacilli and as multicellular aggregates, formed biofilm-like structures embedded in extracellular material along ventricular and inner brain surfaces, bearing pili-like fibers and tubular outer-membrane-derived protrusions. The measured extracellular fibers closely matched the 15-17 nm diameter reported for the smooth chromatin fibers forming neutrophil extracellular traps (NETs) by SEM [30]. NET formation has previously been demonstrated in pneumococcal meningitis, where extracellular DNA networks trapped bacteria but were associated with impaired bacterial clearance [31]. By comparison, the organized fibrous matrix of in vitro *E. coli* biofilms has been shown to primarily depend on curli and cellulose, and extracellular DNA might not be a major structural component [10]. Together with the presence of neutrophils, which we demonstrated by both SEM and histology, this agreement suggests that NET-derived chromatin contributes to the fibrillar scaffold surrounding the bacterial aggregates in E. coli meningoencephalitis, while also considering that other bacterial and host-derived components may be incorporated into a composite extracellular matrix.

Beyond the bacteria-immune system morphological organization, the detected molecules pointed to strong host pressure on bacterial iron acquisition. In nutritional immunity process [11], the neutrophil Lcn-2, detected here only in the infected tissue and CSF, is known to sequester ferric enterobactin and block its use, which is prevented by enterobactin structural modification to glucosylated salmochelins [13]. This well fits the siderophore profile we determined, which showed the absence of enterobactin and a dominant presence of aerobactin- and salmochelin-derived metabolites. DHBS itself is a functional siderophore that delivers iron through the catecholate transporters Fiu, Cir and FepA [32]. Its accumulation suggests a compartment in which the bacteria continue to acquire iron through routes that are poorly captured by Lcn-2, mirroring the catecholate breakdown and aerobactin use we described elsewhere [33]. Calprotectin binds iron [34], zinc and manganese [35]. Its proteoforms converged around ventricular and periventricular regions. Using MALDI MSI, spatial organization of siderophores and calprotectin was previously displayed solely in *Staphylococcus aureus* kidney abscesses[35, 36], where calprotectin, predominantly the S100A8 subunit, formed a rim that encloses the bacterial microcolony. S100A8 and S100A9 predominantly function as a tight heterocomplex. In viral infection, S100A9 homodimer was resolved to be preferentially secreted during inflammation [37], and its activity was mediated by activation of the TLR4-MyD88 pathway, thus regulating the outcome by exaggerating pro-inflammatory response and cell-death [38]. In our model, S100A9-ac and S100A9-ac/m were detected, which suggests modification-specific regulation of S100A9 within the CNS. A second, neutrophil-associated layer, marked by RatNPs, defensin-like peptides, formed ventricular, limbic, and meningeal antimicrobial zones and correlated more closely with calprotectin proteoforms than with siderophores. RatNPs belong to α-defensins stored in neutrophil azurophilic granules [39], and are not produced by resident CNS cells [40]. This supports our findings that both calprotectin and RatNPs co-released with NETs [41]. Contrary to the RatNPs’ antimicrobial and immunomodulatory role, at high local abundance, they can promote vascular dysfunction, and exacerbate neurodegenerative cascades [42]. Counterintuitively, Liao et al. [43] showed, that α-defensins can promote bacterial biofilm formation by targeting the outer membrane protein A of *Acinetobacter baumannii*, including within CSF. Therefore, further studies are needed to exploit the role and outcome of RatNPs in menongoencephalitis.

The next result shifts the interpretation from antimicrobial defense towards neurochemical change. We found that the PEnk-derived peptides globally declined in basal ganglia structures, including regions not dominated by bacterial siderophore signals. A systemic decline of PEnk abundance has been reported in patients with acute neuroinflammation [44], and in neurodegenerative diseases, particularly in Huntington’s and Parkison’s disease [45]. We hypothesize that the route by which the PEnk is affected is indirect. Neuroinflammation propagates from the ventricular, meningeal and perivascular interfaces, where the host-pathogen response was concentrated, into the connected basal ganglia circuits. Such a route is consistent with the reciprocal relationship between opioid tone and neuroinflammation, since δ-opioid signaling restrains microglial pro-inflammatory activation [46], so a fall in enkephalin tone and microglial activation could reinforce one another. To test this, transcript-level or receptor analysis would be required. Nevertheless, our findings add a spatial, neurochemical dimension to meningitis pathology, showing that endogenous neuropeptide circuits are perturbed in parallel with the inflammation, BBB disruption, and edema that typically characterize bacterial meningitis. [24].

CSF provided a translational counterpart to the tissue maps. CSF is the proximal diagnostic fluid for CNS infection. Established CSF markers of bacterial meningitis are bacterial culture and immune proteins such as Lcn-2 that discriminate bacterial from viral meningitis [47], as well as CSF metabolomics that can distinguish bacterial meningitis or CNS infection states [48]. In our model, CSF recovered the same two axes resolved in infected tissue. It raises the prospect of indicating the presence, and potentially the identity, of the organism alongside the magnitude of the antimicrobial response. We used a controlled model with limited sample numbers, therefore validation in etiologically confirmed clinical meningitis will be required before any diagnostic claim can be made.

As with any experimental model, these findings should be read within defined boundaries. To generate a reproducible, anatomically confined infection suitable for high-resolution spatial mapping, we used direct bacteria intracerebral inoculation which consequently bypassed bloodstream invasion and BBB crossing. The single 24 h endpoint captured the spatial organization of the response without considering a temporal sequence. Additional analyses that would define chemical composition of the biofilm matrix and a causal relationship between the mapped species might be beneficial to support our SEM and MALDI MSI findings. These boundaries do not diminish the spatial host-pathogen organization documented here, but define the next experiments, including time-course imaging, matrix-specific validation of bacterial aggregates, and targeted CSF assays in clinically indicated samples.

Overall, we showed that the infected brain behaves as a connected landscape of host-pathogen niches. In CNS interfaces, bacteria compete for iron and form biofilm-like communities. The host responds with Lcn2 and calprotectin ensuring metal sequestration, and forms RatNP-rich antimicrobial zones. In parallel, endogenous neuropeptide circuits, here evaluated at the proenkephalin level, show a decline in abundance. Bacterial localization, metal acquisition, antimicrobial peptides and neuropeptide reduction form the separate but connected maps. The strongest implication demonstrates that CNS infection is defined not only by where bacteria are found, but by where the host response goes.

## Conclusion

Using multimodal imaging, this study reveals experimental E. coli meningoencephalitis as a spatially organized host-pathogen process, in which bacteria and host effector molecules occupy distinct but connected CNS niches. This molecular signature can also be recovered in clinically relevant specimens, e.g., CSF. We provide a comprehensive framework for future host-pathogen molecular interaction studies, while also offering a rationale for mechanistic research into neurochemical infection-induced alterations that may lead to the development of novel or combination therapeutic and diagnostic strategies.

## Experimental Methods

### Materials

9-aminoacridine (9-AA), N-(1-naphthyl) ethylenediamine dihydrochloride (NEDC), 2,5-dihydroxibenzoic acid (DHB), sinapinic acid (SA), staining solutions Hematoxylin Solution Gill No. 3, Differentiation Solution, and Scott’s Tap Water were purchased form Sigma-Aldrich (Prague, Czech Republic). Acetonitrile, methanol, isopropanol, water (LC-MS-grade), chloroform, ethanol, formic acid (FA), trifluoroacetic acid (TFA), acetic acid, and citric acid were purchased from VWR (Stříbrná Skalice, Czech Republic). Standards of enterobactin, aerobactin, ferrioxamine E (FoxE) were purchased from EMC Microcollections (Tübingen, Germany). A standard of mouse Hepcidin 25 was purchased from (Sandhausen Germany). A standard of human neutrophil peptide-1 (HNP-1) was purchased from MedChemExpress (Monmouth Junction, NJ, USA). Luria-Bertani (LB) broth Miller was purchased from HiMedia (Brno, Czech Republic). LB agar used for bacterial cultivation was prepared from agar (15 g/L), NaCl (10 g/L), yeast extract (5 g/L), and tryptone (10 g/L). Phosphate-buffered saline (PBS, pH 7.4) and saline (0.9% NaCl) were prepared from NaCl (8.0 g/L), KCl (0.2 g/L), KH₂PO₄ (0.27 g/L) purchased from Lach-Ner (Neratovice, Czech Republic), and Na₂HPO₄·2H₂O (1.78 g/L) purchased from Honeywell (Seelze, Germany). Sterile PBS and physiological saline were prepared using analytical-grade reagents and ultrapure water.

### Experimental Design

The experimental design considered both animal model development and MALDI MSI methodology. The animal model employed a relatively low bacterial dose while maintaining immunocompetence, allowing progressive bacterial proliferation together with activation and recruitment of immune cells to the brain. This approach mimicked CNS infection and inflammation, enabling spatial investigation of host-pathogen interactions. Cryosectioning was designed so that cultivation, qPCR, histological evaluation, and MALDI MSI analyses were performed from adjacent brain areas to maximize comparability between datasets. Coronal levels at +2.28 and −1.08 mm relative to bregma were selected based on pilot experiments to capture infection-affected brain regions and regions with high endogenous peptide abundance. Infected and control samples were analyzed within the same MALDI MSI analysis to minimize technical variability arising from sample preparation and data acquisition. Prespecified analyses included regional correlation analyses between bacterial and host-derived molecular markers and assessment of their spatial dissociation across brain regions.

### Animals

The studies were performed using three-months-old female LEWIS rats (Charles River, Sulzfeld, Germany). The animals were acclimatized to laboratory conditions for 10 days prior to experimental use and housed under standard laboratory conditions (i.e. 12h light/12h dark day cycle, humidity of 30% to 70%, and temperature of 19 °C to 25 °C) on no-dust bedding in individually ventilated cages with free access to animal chow and water. During the experiments, general health and body weight of the animals were monitored. The number of animals was reduced as much as possible (*n* ≤ 7/group) for all in vivo experiments.

### Experimental induction of *E. coli* neuroinfection

The rats were anesthetized with ketamine (Narkamon; Bioveta, Ivanovice na Hane, Czech Republic) (80 mg/kg, i.p.) and xylazine (Rometar; Bioveta, Ivanovice na Hane, Czech Republic) (10 mg/kg, i.p.) to minimize animal suffering and to prevent animal motion during the surgery. The rats’ heads were shaved off hair. The animals were fixed on a stereotaxic apparatus using ear bars, a snout clamp, and a dental bar, ensuring the precise animal head position for stereotactic surgery. Thermoregulation of the rats during surgery was provided by a heating pad (RightTemp®; Kent Scientific Corporation, Torrington, CT, USA) set at 37 °C. To prevent drying of the cornea, sterile eye ointment (Ophthalmo-Septonex; Zentiva, Prague, Czech Republic) was applied to the eyeballs, and the shaved part of the head was cleaned and disinfected with 4% povidone solution with iodine (Jodisol; SpofaDental, Jicin, Czech Republic). Subsequently, a 2 cm skin incision was made on the scalp, extending from the lambda to just between the eyes, and all soft tissue was removed from the surface of the skull. Using a guide cannula, the site of application was estimated using selected coordinates: Bregma - dx. ML + 1.34 mm; AP + 1.0 mm; DV - 3.0 mm. The application site was marked and then a hole was drilled almost across the width of the skull using a laboratory microdrill (Micro motor handpiece; Saeshin Precision, Daegu, Korea). 5 µL of inoculum (*E. coli* CCM 5172 in the dose of 5˟10^2^ [*n =*7 animals] or 5x10^3^ CFU [*n =*7 animals]) or saline (control group, *n =* 6 animals) was slowly injected at a rate of ˂0.2 µL/s into the prepared hole using a precision 10 µL Hamilton syringe (Hamilton Central Europe, Timisoara, Romania). After application, the drilled hole was covered with sterile surgical hemostatic bone wax (SMI, St. Vith, Belgium) and the incision was sutured with nonabsorbable USP 3/0 - EP2 Polyamide-6 suture (Silon monofilament blue; Chirana T. Injecta, Prague, Czech Republic). Experimental animals were sacrificed under general anesthesia by exsanguination 24 h after inoculation or saline injection. CSF and brain tissue were collected before or right after sacrifice for subsequent analysis. Brain tissues were snap-frozen in ice-cold isopentane. Collected specimens were stored at -80 °C until further use.

### Cryosectioning

The cryosectioning was performed using a Leica CM1950 cryostat (Leica Microsystems GmbH, Wetzlar, Germany). *E. coli*-infected (*n =*5) and control (*n =*4) brain tissues from the animal groups were randomly selected for spatial morphological and molecular analyses. The reduced number of animals was implemented in accordance with the 3Rs principles to minimize animal use while optimizing imaging acquisition time, data quality, and technical reproducibility (Cohen’s d = 2.3, statistical power effect = 84% at the probability level of α = 0.05, determined based on qPCR-data). Frozen brains were transferred to the cryo-chamber and left to temperate for 40 min at -20 °C. The consecutive coronal brain tissue sections from two brain levels, e.g. +2.28 mm and -1.08 mm relative to bregma determined according to the rat brain atlas [49], were collected. For qPCR and cultivation, trimmed 30-µm-thick tissue sections were collected to pre-weighted and sterile Eppendorf tube. Serial 12-µm-thick sections following the previous ones were thaw-mounted on pre-cooled indium tin oxide-coated (ITO) glass slides (Bruker Daltonics, Brno, Czech Republic) and standard glass slides for MALDI MSI and histology, respectively. Brain tissues from either infected or control group were mounted side by side on the same slide to minimize preparation and acquisition variations. All samples were stored at -80 °C.

### Histology

Tissue sections from infected (*n =*5) and control (*n =*4) brains were dried for 40 min and subjected to Gram, and HaE staining to inspect bacterial dissemination and morphological changes, respectively. To demonstrate bacteria, tissues were stained with crystal violet for 5 min, and the excess stain was removed by tap water wash. The stain was stabilized by iodine solution for 2 min and then decolorized for 30 s. A short tap water wash was performed before and after decolorization. Tissues were secondary stained with saphranin for 1.6 min and fixed by 96% and absolute ethanol for 3 min each.

Morphological tissue evaluation was performed upon HaE staining as follows. Tissues were fixed in 95% ethanol for 15 min, stained with hematoxylin for 2.25 min, decolorized with differentiation solution for 40 s, and blue using Scott’s solution for 30 s. A 30 s rinse in running tap water was performed between each step. Tissues were counterstained with eosin for 1 min, and excess stain was removed in tap water for 3 min. Tissues were additionally fixed in absolute ethanol for 1 min. Before permanent mounting using DPX mounting medium (Sigma-Aldrich, Prague, Czech Republic), tissues were cleared with xylene for 15 min. Histological evaluation was performed using a Leica DM2500 microscope (Leica Microsystems GmbH, Wetzlar, Germany). Images were visualized and captured in the LAS X software (Leica Microsystems GmbH, Welzlar, Germany) without further processing. Histological severity was scored from 0 to 4, indicating presence of neuropathological features at 0 = absent, 1 = minimal, 2 = mild, 3 = moderate, 4 = severe level.

### Float-fixation procedure of cryostat sections for scanning electron microscopy analysis

Infected (*n* = 2) and control (*n* = 2) brain tissues were prepared for the SEM analysis according to the published protocol [29]. Briefly, 1.5 mL of fixative solution (3% glutaraldehyde, 0.1 M cacodylate buffer) was added into a six-well plate and frozen at -20 °C. The plate was conditioned with the tissues in a cryo-chamber for 40 min. 70 µm-thick sagittal tissue sections cryosectioned using the Leica CM190 cryostat were transferred onto a frozen fixative surface, and the plate was placed in a refrigerator. The tissue sections were gently fixed by slowly melting the fixative overnight, followed by tissue floating wash on the cacodylate buffer level for 20 min, three times. Washed sections were mounted on coverslips and packed in fine copper wire mesh. Subsequently, the sections were dehydrated in a graded ethanol series (25, 50, 75, 90, 96, 100, and 100%) and dried in a critical point dryer K850 (Quorum Technologies Ltd., Laughton, UK). CPD-dried sections were mounted onto aluminum specimen stubs using conductive tape and sputter-coated with 3 nm of platinum in a high-resolution Turbo-Pumped Sputter Coater Q150T (Quorum Technologies Ltd., Ringmer, UK).

### Scanning electron microscopy

Prepared tissue samples were examined in a FEI Nova NanoSEM 450 scanning electron microscope (Thermo Fisher Scientific, Brno, Czech Republic) at acceleration voltage 5 kV and spot size 3 using Everhart-Thornley Detector (ETD), Circular Backscatter Detector (CBS), and Through the Lens Detector (TLD). The original 16-bit images 1536 × 1103 px were recorded at primary magnifications ranging from thousands to hundreds of thousands times. For presentation, the gamma of the original 16-bit images was adjusted if necessary, and the images were converted to 8-bit ones without any other processing.

### Quantitative PCR analysis

Coronal brain sections (30 μm) from infected (*n* = 5) and control (*n* = 4) brain consecutive to those analyzed by MALDI MSI were lyophilized for 48 h, homogenized, and 5 mg of dry tissue was used for DNA extraction with the DNeasy Blood & Tissue Kit (Qiagen, Germany). DNA concentration was determined using a Qubit 2.0 fluorometer and the Qubit dsDNA HS Assay Kit (Thermo Fisher Scientific, USA). Primers and a hydrolysis probe targeting the *E. coli uidA* gene were designed using PrimerQuest and verified by BLAST analysis (Table S9). qPCR was performed using iTaq Universal Probes Supermix (Bio-Rad, USA) on a CFX96 Real-Time PCR System (Bio-Rad, USA). Reactions (20 μL) contained 300 nM of each primer, 250 nM probe, and 5 μL template DNA. Cycling conditions were 95 °C for 5 min, followed by 40 cycles of 95 °C for 5 s and 60 °C for 30 s. Samples and standards were analyzed in technical triplicates. A six-point standard curve (10^2^-10^7^ CFU/mL [data not shown]) was generated using homogenized noninfected rat brain tissue spiked with *E. coli*. No-template controls contained PCR-grade water. Bacterial abundance was calculated from Cq values using CFX Manager v3.1 (Bio-Rad) and expressed as genome equivalents.

### Bacterial culture and CFU determination

For bacterial CFU determination, collected coronal 30-μm brain sections from infected (*n* = 5) and control (*n* = 4) tissues at both +2.28 mm and -1.08 mm level from bregma and consecutive to those used for qPCR were suspended in 600 μL PBS, vortex-homogenized, and plated on LB agar. Samples with high bacterial loads were additionally analyzed at a 10^-1^ dilution. All cultures were plated in technical triplicates, incubated at 37 °C for 24 h, and quantified by colony counting.

### Sample preparation for MALDI qMSI

Three different sample preparation protocols were applied to analyze siderophores, antimicrobial peptides, and intact proteins in infected (*n =* 5) and control (*n* = 4) tissue sections. On the measurement day, tissue sections were desiccated for 40 min and scanned using the flatbed optical scanner (Epson Perfection V600 Photo, Nagano, Japan).

Siderophores were imaged and quantified [50] in brain tissues using a sample preparation protocol as follows. Calibration standard solutions of a ferri-aerobactin and desferri-enterobactin were prepared at the concentration of 10, 25, 50, 100, 250, 500, 1000, 5000, 10000, and 5000 µg/mL, including blank solution and quality control samples of 1000 (low level) and 2500 (high level) µg/mL in 50% methanol. The calibration solutions (0.3 µL) were manually spotted on the control tissue section in six replicates. Infected, control sections, and sections with the spotted calibration curve were then sprayed with a matrix mixture of 9-AA and NEDC with a concentration ratio of 1:2, dissolved in 75/20/5% of methanol/acetonitrile/water using a SunCollect sprayer (SunChrom GmbH, Friedrichsdorf, Germany). The following matrix spraying conditions were applied: sprayer tip position in Z axis with Z offset of 30 and 5 mm, respectively, and 13 layers of the matrix sprayed horizontally with a line distance of 2 mm, a constant flow rate of 70 µL/min, nitrogen gas pressure of 2.1 bar, and sprayer head speed of 900 and 1700 mm/min in X and Y axis, respectively.

Antimicrobial peptides were visualized and semi-quantified upon application of a modified previously published protocol [50, 51]. Briefly, calibration standard solutions of HNP-1 with a concentration of 50, 100, 250, 500, 750, 1000, 2500, 5000, 7500, and 10000 µg/mL were prepared in 50% methanol, including blank and quality control samples of 1000 (low level) and 2500 (high level) µg/mL. The same abovementioned approach was used for spotting the HNP-1 calibration curves. Prior to MALDI matrix application, the internal standard of mouse hepcidin 25 (10000 µg/mL in 40% ethanol/1% TFA) was homogenously sprayed over the infected and control sections, and section with calibration standars using the SunCollect sprayer with the following conditions: the sprayer head position in Z axis of 30 mm with a Z offset of 5 mm; 10 layers of matrix sprayed horizontally with a distance between lines of 2 mm, increasing flow rate of 20, 35, and 50 µL/min for the first three layers and constant 70 µL/min for the rest at the constant nitrogen pressure of 2.1 bar, and sprayer head speed of 900 and 1700 mm/min in X and Y axis, respectively. After 20 min of vacuum-drying, brain lipids were washed from the tissues by a 35-second chloroform wash, followed by a quick dry with a stream of nitrogen and an additional 20 min of vacuum-drying. The sample preparation was completed by applying a DHB matrix with a concentration of 25 mg/mL dissolved in 40% ethanol/1% TFA using the same spraying conditions as for the internal standard solution.

Intact proteins were visualized and relatively quantified based on a published protocol [52] with modifications. Tissue sections were subjected to a series of ethanol submersions: 70%, 90%, 95%, and absolute ethanol for 30 s each, followed by a wash in the Carnoy’s solution containing chloroform, methanol, and acetic acid (6:3:1 v/v) for 3 min. Tissues were then submerged in absolute ethanol, pure water, and again absolute ethanol for 30 s each. Before applying a MALDI matrix, tissues were dessicated for 15 min. Sinapinic acid (10 mg/mL dissolved in 70% ethanol/1% TFA) was sprayed over the tissues using a M3+ Sprayer (HTX Technologies, LLC, Carrboro, North Carolina, USA) with the following parameters: nozzle temperature of 80 °C, 8 passes with a track spacing of 1.5 mm, a constant flow rate of 100 µL/min and nitrogen gas pressure of 10 psi, and cross spraying pattern. Tissues were rehydrated, and the matrix was recrystallized by suspending prepared samples over 1 mL of 5% acetic acid for 3.5 min at 85 °C, followed by 5 min of drying at room temperature.

### MALDI MSI data acquisition

MALDI MSI analyses of siderophores and peptides were performed using the solariX 12T-2ω Fourier transform ion cyclotron resonance (FTICR) mass spectrometer equipped with a Smartbeam II 2 kHz laser (Bruker Daltonics, Billerica, USA).

MSI data of siderophores were acquired in negative ion mode using quadrature phase detection in a mass range from 150 to 1500 *m/z* externally calibrated against clusters of red phosphorus. The ion signal at *m/z* 885.549853 (phosphatidylinositole 38:4, [M-H]^-^, a lipid naturally occurring in the brain tissue) was used for online data calibration. Data was collected with 2 M data points in the transient with 0.4 s transient length providing an estimated resolving power of 190,000 at *m/z* 400. Ion source was optimized to funnel 1 (-150 V), skimmer 1 (-15 V), and funnel amplitude (120 Vpp). Ion optic was optimized to collision cell voltage (CV, +3.0 V; DC bias: -1.5 V), time-of-flight delay (TOF, 0.7 ms), and transfer optics (4 MHz, Q1 *m/z* 200). The laser power was tuned at the beginning of the experiment, and the parameters were kept constant throughout each MALDI MSI data acquisition (100 laser shots/position). All MSI data were acquired with 100 μm lateral resolution, and 98% data reduction.

MSI data of peptides were collected in positive ion mode using the quadrature phase detection. The acquisition method was calibrated on the clusters of red phosphorus prior to analysis. The ion signal at *m/z* 2755.026519 (the third isotope of the mouse hepcidin 25, [M+H]^+^) was utilized for internal calibration. Spectra were obtained by collecting data from *m/z* 300 to 4000 at one spectrum per pixel. The time domain file size was set at 1 M, with an estimated resolving power of 190,000 at *m/z* 400. The ion optics were optimized as follows: the source optics deflector plate (220 V), funnel 1 (150 V), and amplitude of 200 Vpp; the transfer optics CV (-2.5 V), DC bias (0.8 V), collision RF frequency (2 MHz) with an amplitude of 1200 Vpp, and TOF delay (1.3 ms), frequency (2 MHz) and Q1 at *m/z* 300. The laser operated at a frequency of 1000 Hz (100 shots/position) with the laser power tuned before data acquisition and maintained constant throughout the analysis. Data were collected using 100-μm raster steps, and 98% data reduction.

MALDI MSI analysis of intact proteins was performed using the Ultraflex II MALDI-TOF/TOF mass spectrometer (Bruker Daltonics, Billerica, USA). MSI data were acquired in a positive linear ion mode in a mass range from 3,000 to 30,000 *m/z* externally calibrated against clusters of red phosphorus and ProtMix I and II standards (Bruker Daltonics, Bremen, Germany). Instrumental conditions were tuned to high voltage of ion source 1 (25 kV), ion source II (23.45 kV) and lens (6.05 kV), and pulsed ion excitation (90 ns). The laser power was tuned before data acquisition and kept constant throughout the analysis. Data were collected with a raser step of 100µm, by summing 100 laser shots per position.

### MALDI MSI data processing and validation

All MSI data were visualized and processed in SCiLS Lab v.2026b Pro software (both Bruker Daltonics, Bremen, Germany). Siderophore, peptide, and protein data were processed separately, and a SCiLS dataset included both infected and control experimental group, and calibration curves if applicable. Import parameters were as follows centroid (siderophores and peptides) and reduced (proteins) spectra; axis bin size at minimum-maximum *m/z* (siderophores, 0.03-3.1 mDa; peptides, 0.2-73 mDa, and proteins, 3.4 mDa) and narrow baseline removal for proteins. The resulting number of data points were 3846502, 362944, and 7947 for siderophores, peptides, and proteins, respectively. Data of siderophores, peptides, and proteins were normalized to root-mean-square, against the peak area of internal standard of mouse hepcidin 25, and to total ion count, respectively. Regions of interest (ROI) were manually drawn according to the rat brain atlas [49] using the optical image and the fiducial markers present in the brain tissue. The normalized average peak areas within ROI, considering either deprotonated (for siderophores) or protonated, sodiated and potassiated (for peptides) ion species, and average intensity of peak maximum for protonated proteoform ions were used for data extrapolation. For quantitative data assessment, the density of rat brain (1.027 g/cm^3^), and the thickness of tissue section (12 µm) were considered. Data were finally expressed as mean ± standard deviation (SD). For matching ions of bacterial dissemination and immune response, spectra were exported from SCiLS software and imported into mMass software [53], and detected pekas were primarily identified by database search (www.hmdb.ca, [54] www.uniprot.org [55], in-house build siderophore database, and previously published MSI-detected peptide list [56]) based on the mass accuracy with a mass error below 5 ppm, and isotopic profile provided by high-resolution MS analyses for siderophores and peptides, and a mass error below 100 ppm for proteins in addition to compound distribution comparison with published data [56, 57]. UMAP was used for nonlinear dimensionality reduction of MALDI MSI peptidomic data before unsupervised spatial clustering. For image analysis intensities, compounds of interest were displayed at a gray scale, and images (8-bit depth, resulting in signal levels from 0 to 250) were exported as PNG files into the ImageJ software. Intensities were reported as surface spanning from left to right brain hemisphere with a depicted line of 7-10-pixel thickness. Intensity profile data were exported as .csv files and processed in the GraphPadPrism 10.0.1 for Windows (GraphPad Software, San Diego, California, USA) [53].

### Validation of MALDI MSI methods

The MALDI MSI methods were validated for linearity, coefficient of determination (R2), limit of detection (LOD), limit of quantification (LOQ), within-run accuracy, precision, selectivity, and specificity. The validation followed the guidelines established by U.S. regulatory agencies for the validation of bioanalytical methods (https://www.fda.gov/regulatory-information/search-fda-guidance-documents/bioanalytical-method-validation-guidance-industry). For MSI data, validation requirements were adapted as published elsewhere.[50] For MS and MSI data, calculation of LOD and LOQ was performed via the LINEST function, using the following expressions respectively: LOD=3.3x σ/S and LOQ=10x σ/S (S, slope of the calibration curve; σ, SD of the y-intercept). The strength of the linear response was assessed using Pearson’s correlation coefficient *r*. Specificity was confirmed by the MS instrument’s resolving power, achieving mass resolution at compound-specific *m/z* value while detecting no other ion signals. The validation parameters are reported in Table S2 and Figure S2.

### LC-MS analysis of siderophores in CSF and brain tissue homogenate

Siderophores were semi-quantified by LC-MS in collected infected (*n* = 5) and control (*n* = 4) CSF and brain samples. CSFs (10 µL) were spiked with an internal standard FoxE (5 µL, 1 µg/mL in 50% acetonitrile). Brain homogenates were weighed (7–12 mg) into 2.0 mL vials, dissolved in water (200 µL), and thoroughly ultrasonicated. A portion (100 µL) of the homogenate solutions were spiked with FoxE (10 µL, 2 µg/mL in 50% ACN). Siderophores were extracted from the CSF and brain homogenate samples by pre-cooled isopropanole (400 µL). After 1 h in -80 °C, samples were centrifuged for 10 min at 4 ℃ and 14 000 g, and supernatants were transferred into a new vial and concentrated to dryness for 2 h at 35 ℃. Before LC-MS, samples were resuspended in 5% acetonitrile. Samples were analyzed using a Dionex UltiMate 3000 UHPLC liquid chromatography system (Thermo Fisher Scientific, Massachusetts, USA) connected to a solariX 12T-2ω FTICR mass spectrometer. Samples (1 µL) were injected onto an Acquity HSS T3 C18 analytical column (1.8 µm, 1.0 × 150 mm; Waters Corporation, Massachusetts, USA) pre-heated at 40 ℃. Analytes were separated within a 13-min gradient elution of 0.1% formic acid in 1% and 99% acetonitrile with a flow of 50 µL/min as follows: 2% acetonitrile (0-1 min), 99% acetonitrile (1-9 min), 99% acetonitrile (9-10 min), 2% acetonitrile (10.0-10.1 min), and 2% acetonitrile (10.1-13.0 min). Mass spectra were collected in the electrospray ionization positive ion mode within a mass range from 200 to 1,200 *m/z*. The mass spectrometer was externally calibrated against clusters of 0.01% sodium trifluoroacetate. In addition, MS spectra were online calibrated against standard mixture (ESI Tuning Mix for Ion Trap, Sigma-Aldrich, Prague, Czech Republic). The system sustainability test of the LC-MS system was regularly checked by analyzing the HPLC peptide standard mixture (Sigma-Aldrich, Prague, Czech Republic). The MS method parameters were optimized as follows. Data were collected with 256 k data points in the transient with 0.14 s transient length with an estimated resolving power of 33,000 at *m/z* 400 and ion accumulation time of 0.25 s. Ion source was optimized to capillary exit (200 V), funnel 1 (-150 V), skimmer 1 (-15 V), and funnel amplitude (150 Vpp). The electrospray parameters were optimized to capillary voltage (-4500 V), end plate offset (-600 V), nebulizer N2 gas (0.5 bar), dry N2 gas (2.5 L/min), and dry gas temperature (240 °C). Ion optics were optimized to CV (-10 V); DC bias: (+1.0 V), TOF (0.9 ms), and transfer optics (4 MHz). All MS data were acquired with 98% data reduction. Identification of siderophores was confirmed by matching retention time, isotope pattern, and product ion mass spectra generated by MS/MS using a collision-induced dissociation. For MS/MS analysis, the ions were isolated with a 10 Da mass width and dissociated with collision energy of 0.4, and 8 V. Obtained data were processed and evaluated with the DataAnalysis 6.0 (Bruker Daltonics, Bremen, Germany), CycloBranch 2.1.35. Using extracted ion chromatograms with a 0.005-Da spectral width, the responses were summed from the integrated peak areas of ferri- and desferri-ion species and normalized to the response of an internal standard. CSF and brain homogenate samples were measured in technical duplicates. Data were expressed as mean ± SD. In the brain homogenates, data was recalculated to the weight of the corresponding brain tissue.

### MALDI MS analysis of antimicrobial peptides in CSF

Antimicrobial peptides were semi-quantified in crude samples of infected (*n* = 5) and control (*n* = 4) CSFs by MALDI MS. Infected and control samples were diluted three times with a solution of mouse hepcidin 25 internal standard with a concentration of 500 µg/mL prepared in 50% acetonitrile. 0.5 µL of each sample was spotted on the MALDI ground steel target in five replicated and overlaid with 0.5 µL of α-cyano-4-hydroxycinnamic acid matrix (10 mg/mL in 50% acetonitrile/0.2% TFA). Data from each spot were averaged from 32 scans acquired in a random pattern with 300 laser shots/scan. MS data were acquired using solariX 12T-2ω FTICR mass spectrometer in positive ion mode in a mass range from 300 to 4000 *m/z.* with an external calibration against clusters of red phosphorus. The MS method parameters were optimized as follows. Data was collected with 1 M data points in the transient with 0.8 s transient length with an estimated resolving power of 190,000 at *m/z* 400. Ion source was optimized to funnel 1 (-150 V), skimmer 1 (-15 V), and funnel amplitude (120 Vpp). Ion optics were optimized to CV (-1.5 V); DC bias: +1.0 V), TOF (2 ms), and transfer optics (2 MHz). The laser power was kept constant during data acquisition. All MALDI MS data were acquired without data reduction. MALDI MS data were processed in DataAnalysis 6.0 (Bruker Daltonics, Bremen, Germany). Absolute intensities of protonated, sodiated and potassiated ion species of compound of interest were summed and normalized to the absolute intensity of the internal standard. Data were expressed as mean ± SD. All controls were negative to RatNPs.

### Analysis of rat lipocalin-2 in infected and control brain homogenates and cerebrospinal fluid

Quantitative analysis of rat lipocalin-2 (Lcn-2) in cerebrospinal fluid (CSF; infected, *n* = 5 animals; control, *n* = 8 animals) and brain homogenates (infected, *n* = 5 brains; control, *n* = 4 brains) was performed using the RayBio® Rat Lipocalin-2/NGAL ELISA Kit (RayBiotech, GA, USA) according to the manufacturer’s instructions. Lyophilized brain tissue (5 mg dry weight) was extracted in 25% acetonitrile containing 1% formic acid using probe sonication for 10 s. Brain homogenate extracts and CSF samples were serially diluted in the assay diluent to identify dilution factors within the linear range of the calibration curve. Final quantification was performed at 500- or 1000-fold dilution for brain homogenate extracts and at 5000-fold dilution for CSF samples. All samples were then processed according to the manufacturer’s protocol. Absorbance was measured at 450 nm using a SpectraMax Plus 384 Microplate Reader (Molecular Devices, CA, USA). Lcn-2 concentrations were calculated from the calibration curve, considering assay-specific limits of detection and quantification. The limits of detection and quantification were 38 and 130 pg/mL for brain homogenate extracts and 15 and 30 pg/mL for CSF, respectively. For brain homogenates, final values were recalculated and expressed as ng of Lcn-2 per mg of dry brain tissue.

### Statistical analysis

Statistical analyses were performed using GraphPadPrism 10.0.1 for Windows. Descriptive statistics reported the means and SD, standard error of the mean, coefficient of variation, and 95% confidence interval. Data was tested for normality using the Shapiro-Wilk normality test. If the normality test was passed or not, parametric or nonparametric statistical tests, respectively, were used. Statistical analyses were performed on data obtained from infected (*n* = 5) and control (*n* = 4) brain tissues. qPCR and cultivation data were statistically tested using two-tailed Mann-Whitney test. Histological severity score was assessed using Kruskal-Wallis test followed by an uncorrected Dunn’s multiple-comparison test.

Regional MALDI MSI data from infected animals were analyzed separately at +2.28 and -1.08 mm relative to bregma. The correlations between siderophores, proteins, and RatNPs were determined using Sperman correlation analysis. Differences in antimicrobial peptides in infected samples were tested using paired Friedman test followed by Dunn’s multiple comparison. The relative abundance of all determined peptides was compared between control and infected samples using unpaired two-tailed t-test. Differences in abundance of PEnks were evaluated using Welch’s t-test. At individual PEnk peptide level, the differences between control and infected samples were determined by unpaired t test with Welch correction, followed by Bonferroni-Dunn’s multiple comparisons test.

## Acknowledgements

Czech Science Foundation (22-06771S) to DL; Ministry of Education, Youth and Sports of the Czech Republic (CZ.02.01.01/00/22_008/0004597, Talking microbes - understanding microbial inter-actions within One Health framework) to VH, and by National Institute of Virology and Bacteriology (Programme EXCELES, ID project no. LX22NPO5103)−funded by the European Union−Next Generation EU; the Ministry of Education, Youth and Sports of the Czech Republic (project EATRIS-CZ LM2023053 to MP. The authors used services of the Czech Centre for Phenogenomics at the Institute of Molecular Genetics supported by the Czech Academy of Sciences RVO 68378050 and by the project LM2023036 Czech Centre for Phenogenomics provided by Ministry of Education, Youth and Sports of the Czech Republic.

## Author contributions

D.L. designed and conceptualized the study. D.L., T.H.J., O.B., G.L., H.M., L.K., H.M., J.H., K.D.B., M.P., M.P., A.P., and E.B. performed experiments or contributed analytical methods. D.L., T.H.J., O.B., L.K., A.P., J.N. and V.H. analyzed data. D.L. supervised the project and performed formal analysis of all data. D.L. and V.H. wrote the manuscript with input from all authors. All authors reviewed and approved the final manuscript.

## Data Availability Statement

The raw mass spectrometry data can be downloaded from https://doi.org/10.57680/asep.0651117 and viewed using the CycloBranch software, https://ms.biomed.cas.cz/cyclobranch/ of SCiLS lab software. All remaining data are available in the main text or the supplementary materials.

## Ethics Statement

All animal experiments were performed in accordance with the regulations and guidelines of the Czech Animal Protection Act (No. 246/1992), and with the approval of the Czech Ministry of Education, Youth, and Sports (MSMT-8849/2022-4), and the institutional Animal Welfare Committee of the Faculty of Medicine and Dentistry of Palacký University in Olomouc.

