## Supplementary Materials for "Multimodal Imaging Reveals Spatial Host-Pathogen Microenvironments in *Escherichia coli* Meningoencephalitis"

Short title

Spatial *E. coli* Meningoencephalitis

**Authors**

Dominika Luptáková,^1^* Tereza Hřivnová Juříková,^1^ Oldřich Benada,^1^ Gabriela Lokočová,^1^ Helena Marešová,^1^ Hynek Mácha, ^1^ Jiří Houšť, ^1^ Kateřina Dvořáková Bendová,^2^ Miroslav Popper,^2^ Miloš Petřík,^2,3,4^ Andrea Palyzová,^1^ Lukáš Kučera,^5^ Evgeniya Biryukova,^1,6^ Jiří Novák,^1^ Vladimír Havlíček^1,7^*

**Affiliations**

^1^Institute of Microbiology of the Czech Academy of Sciences, Vídeňská 1083, 142 00 Prague, Czech Republic

^2^Institute of Molecular and Translational Medicine, Faculty of Medicine and Dentistry, Palacký University, Hněvotínská 5, 779 00 Olomouc, Czech Republic

^3^Czech Advanced Technology and Research Institute, Palacký University, Šlechtitelů 241/27, 779 00 Olomouc, Czech Republic

^4^Institute of Molecular and Translational Medicine, University Hospital, Hněvotínská 5, 779 00, Olomouc, Czech Republic

^5^Institute of Molecular Genetics, Academy of Sciences of the Czech Republic, Vídeňská 1083, 142 00 Prague, Czech Republic

^6^Department of Biochemistry, Faculty of Science, Charles University, Albertov 6, 128 00 Prague 2, Czech Republic

^7^Department of Analytical Chemistry, Faculty of Science, Palacky University in Olomouc, 17. listopadu 1192/12, 779 00 Olomouc, Czech Republic

***Corresponding authors**

Dominika Luptáková, and Vladimír Havlíček,

**Other Supplementary Materials for this manuscript include the following:**

The raw mass spectrometry data and MALDI MSI processed data can be downloaded from https://doi.org/10.57680/asep.0651117 and viewed using the CycloBranch https://ms.biomed.cas.cz/cyclobranch/ and SCiLS software.

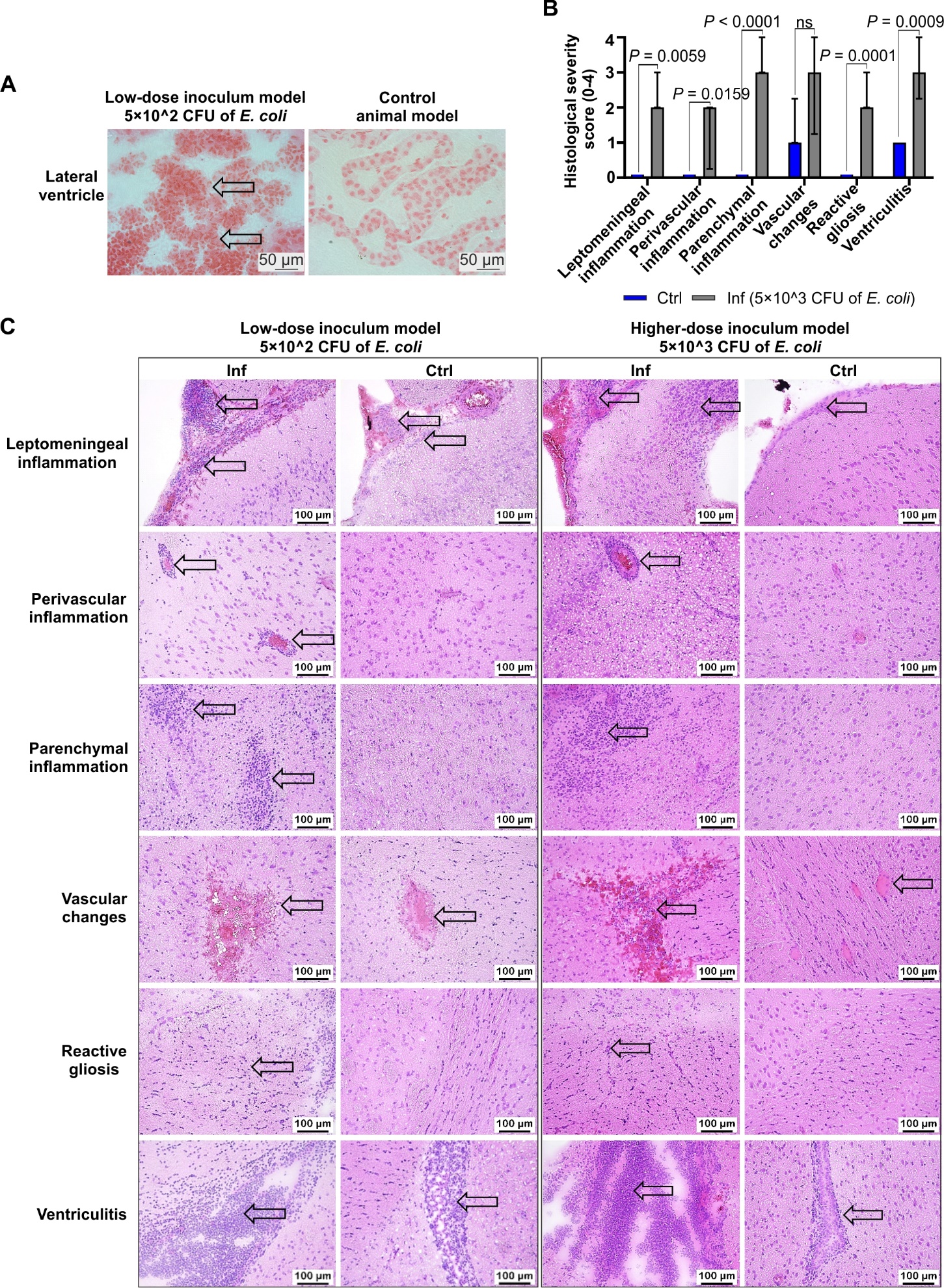

Figure S1. Histopathological analysis of the experimental rat models of *E. coli*-induced meningoencephalitis. (A) Representative images of Gram-stained *E. coli-*infected (inoculation dose of 5˟10^2 CFU of *E. coli*) and control brain tissue section at the level of -1.08 mm from bregma. Gram negative *E. coli* bacteria were typically pink stained (arrow) with high abundance in ventricles. Scale bars, 50 µm. (B) Histopathological scoring (0–4 scale) of CNS pathology in control (ctrl, *n* = 5) and infected (inf, *n* = 5) animals inoculated with 5˟10^3 CFU of *E. coli* at both +2.28 mm and -1.08 levels relative to bregma. Data are presented as median ± interquartile range. Differences in bacterial load and histopathological scores between control and infected groups were assessed using the Kruskal–Wallis test followed by an uncorrected Dunn’s multiple-comparisons test. *P* values are indicated in the graphs. ns, not significant. (C) Representative hematoxylin and eosin-stained tissue sections showing characteristic neuropathological features (arrows) at the brain level of +2.28 mm from bregma from both experimental infection low-dose inoculum model (inoculation dose of 5˟10^2 CFU of *E. coli*, *n* = 5 animals) and higher-dose inoculum model (inoculation dose of 5˟10^3 CFU of *E. coli*, *n* = 5 animals) compared to controls (*n* = 4 and 5 animals for low-dose and higher-dose inoculum, respectively). Scale bars, 100 μm.

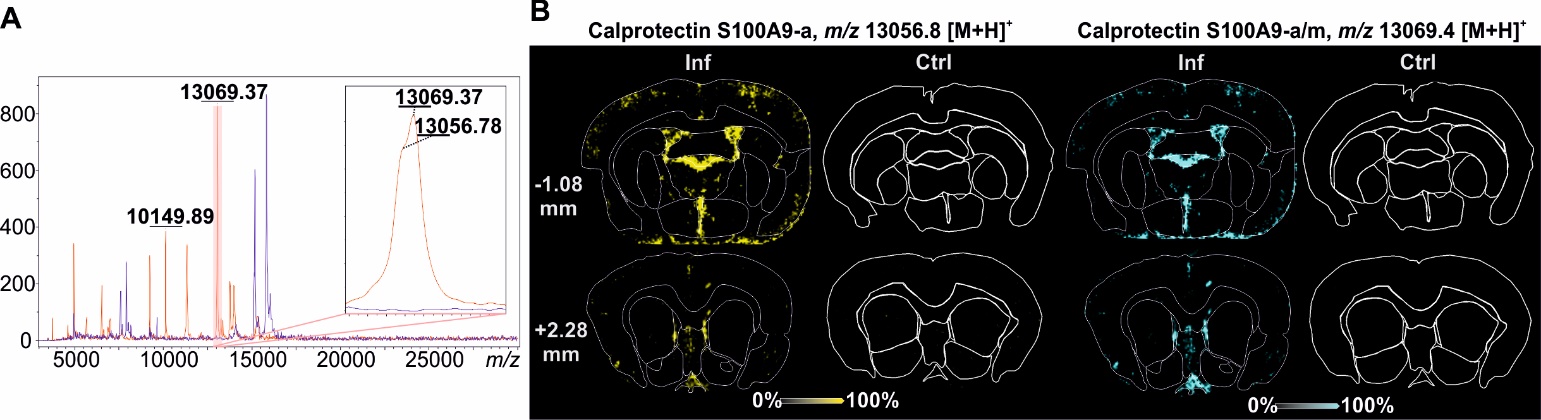

Figure S2. Visualization of calprotectin proteoforms by MALDI TOF/TOF-MSI. (A) Average MALDI mass spectra acquired over the *m/z* 3,000 - 30,000 range from infected (*n* = 5, red) and control (*n* = 4, blue) brain tissues. Peaks corresponding to the calprotectin subunit S100A8 (*m/z* 10,149.89), acetylated S100A9 (S100A9-a, *m/z* 13,056.78), and acetylated/methylated S100A9 (S100A9-a/m, *m/z* 13,069.37) are indicated. The inset shows an expanded view of the S100A9 proteoforms. (B) Representative MALDI MSI ion images of S100A9-a (*m/z* 13,056.8) and S100A9-a/m (*m/z* 13,069.4) in infected (Inf) and control (Ctrl) brain sections at two coronal levels relative to bregma (+2.28 mm and −1.08 mm). MSI data were normalized to the total ion count (TIC). Ion signal intensities are displayed as relative abundance on a 0-100% color-coded scale. TOF/TOF, tandem time-of-flight.

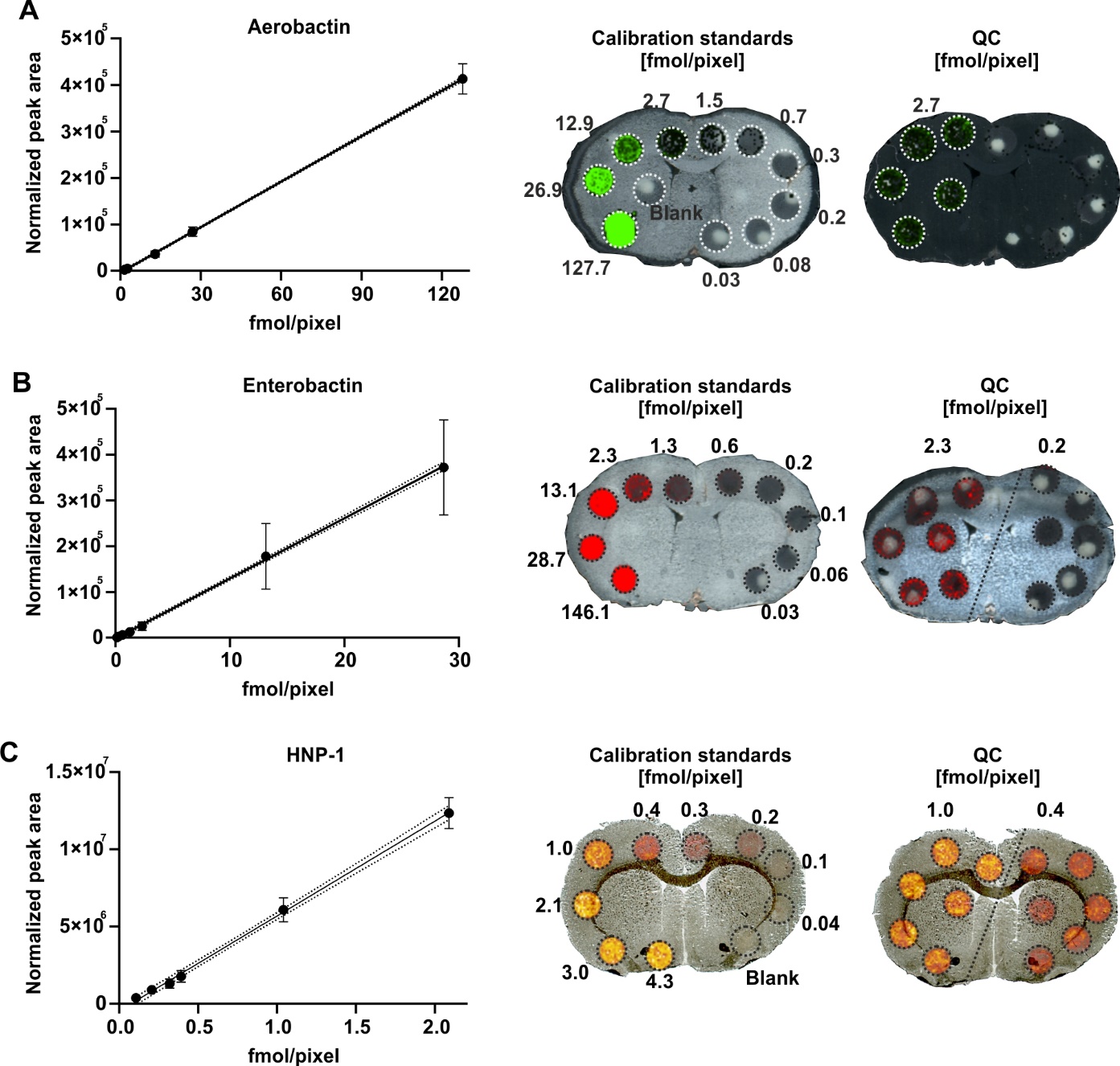

Figure S3. Calibration curves generated by MALDI qMSI. Calibration curves for (A) aerobactin, (B) enterobactin, and (C) human neutrophil peptide-1 (HNP-1) were generated by plotting the amount of analyte per pixel (fmol/pixel) against the root-mean square-normalized average peak area of aerobactin and enterobactin and mouse hepcidin 25-normalized average peak area of HNP-1 measured within each calibration spot. Calibration data were fitted using simple linear regression. Representative MALDI MSI images of calibration standards and quality control (QC) spots are shown. QC spots at independent concentration levels were used to assess method performance and calculate validation parameters, including accuracy and precision as shown in Table S2. MALDI qMSI, matrix-assisted laser desorption/ionization quantitative mass spectrometry imaging.

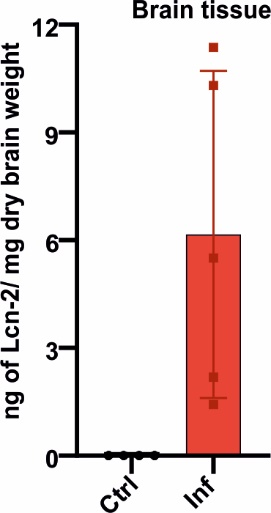

**Figure S4. Determination of lipocalin-2 induction during *E. coli* meningoencephalitis.** Quantitative ELISA analysis of rat lipocalin-2 (Lcn-2) in brain tissue extracts from control (Ctrl, *n* = 4) and *E. coli*-infected (Inf, *n* = 4) animals. Brain tissue values are expressed as ng of Lcn-2 per mg dry brain tissue weight. Bars show mean ± standard deviation.

**Table S1. List of assigned *E. coli* characteristic siderophores with high mass accuracy as determined by MALDI FTICR-MSI experiments.**

| **Siderophore** | **Molecular formula** | **Observed *m/z*,**  **[M-H]^-^** | **Theoretical *m/z*, [M-H]^-^** | **Error ppm** |
| --- | --- | --- | --- | --- |
| **Aerobactin** | C_22_H_36_N_4_O_13_ | 563.2207 | 563.2206 | 0.2 |
| **DHBS** | C_10_H_11_NO_6_ | 240.0516 | 240.0514 | 0.9 |
| **Salmochelin SX** | C_16_H_21_NO_11_ | 402.1047 | 402.1042 | 1.4 |

*E. coli* siderophores were detected in negative ion mode as deprotonated ion species mainly. Additionally, aerobactin was detected in sodiated [M+Na-2H]^-^, pottasiated [M+K-2H]^-^, and dehydrated [M-H_2_O-H]^-^ ion forms.

**Table S2. Validation parameters of the matrix-assisted laser desorption/ionization quantitative mass spectrometry imaging (MALDI qMSI) used to (semi)quantify rat α-defensins and siderophores.**

|  | **QC level [µg/g] of HNP-1/aerobactin/enterobactin** | **Aerobactin** | **Enterobactin** | **HNP-1** |
| --- | --- | --- | --- | --- |
| **Linear regression equation** |  | y = 3265.2x – 4067.7 | y = 13135x – 1930.1 | y = 6143057x – 463268 |
| **Linear calibration range [fmol/pixel]** |  | 1.5 – 127.7 | 0.1 – 28.7 | 0.1 – 2.1 |
| **Coefficient of determination (R^2^)** |  | 0.9999 | 0.9994 | 0.9988 |
| **LOD [µg/g]** |  | 3.9 | 2.4 | 1.7 |
| **LOQ [µg/g]** |  | 11.5 | 7.3 | 5.1 |
| **Recovery [%]** |  | 102 | 106 | 103 |
| **Precision [RSD, %]** | 12.3/n.a./1.3 | n.a. | 8.0 | 9.9 |
|  | 30.9/12.9/13.9 | 10.9 | 20.0 | 9.3 |
| **Accuracy [%]** | 12.3/n.a./1.3 | n.a. | 127 | 102 |
|  | 30.9/12.9/13.9 | 109 | 89 | 89 |
| **Specificity [%]** |  | 100 | 100 | 100 |

The analytical performance characteristics evaluated included linearity, calibration range, coefficient of determination (R²), limit of detection (LOD), and limit of quantification (LOQ), precision, accuracy, and specificity. These parameters were established using calibration standards of human neutrophil peptide-1 (HNP-1), aerobactin and enterobactin at five or six different concentrations, blanks, and quality control samples at up to two concentration levels (12.3 and 30.9 µg/g of brain tissue for HNP-1; 12.9 µg/g of brain tissue for aerobactin; 1.3 and 13.9 µg/g of brain tissue for enterobactin), measured in five or six technical replicates. Peptide and siderophore data were normalized to peak area of mouse hepcidin 25 and root-mean-square, respectively, prior to curve fitting. RSD, relative standard deviation; n.a., not applicable.

**Table S3. List of assigned forms of the rat calprotectin protein as determined by MALDI-TOF/TOF-MSI experiments.**

| **Calprotectin proteoforms** | **Protein sequence** | **PTM** | **Molecular formula** | **Observ. *m/z*, [M+H]^+^** | **Theor. *m/z*, [M+H]^+^** | **Error ppm** | **UniProt ID** |
| --- | --- | --- | --- | --- | --- | --- | --- |
| S100A8 (2-89) | ATELEKALSNVIEVYHNYSGIKGNHHALYRDDFRKMVTTECPQFVQNKNTESLFKELDVNSDNAINFEEFLVLVIRVGVAAHKDSHKE | ac. (1x) | C_450_H_698_N_124_O_140_S_2_ | 10149.9 | 10150.1 | -20 | P50115 |
| S100A9 (2-113) | AAKTGSQLERSISTIINVFHQYSRKYGHPDTLNKAEFKEMVNKDLPNFLKREKRNENLLRDIMEDLDTNQDNQLSFEECMMLMGKLIFACHEKLHENNPRGHDHSHGKGCGK | ac. (1x) | C_562_H_891_N_169_O_174_S_8_ | 13056.8 | 13056.4 | 31 | P50116 |
|  |  | ac. (1x), m. (1x) | C_563_H_921_N_169_O_174_S_8_ | 13069.4 | 13070.4 | -77 |  |

Proteins were detected in positive ion mode in their poronated forms. PTM, posttranslational modification; Observ., observed; Theor., theoretical; ac., acetylation; m., methylation.

**Table S4. Spearman correlation analysis of the detected *E. coli* siderophores DHBS, aerobactin and salmochelin SX with immune system–derived rat antimicrobial peptides (RatNPs) and rat proteoforms of calprotectin (S100A8, S100A9-ac, and S100A9-ac/m) in infected brain tissues (*n* = 5).**

| **Brain level from bregma (mm)** | **Spearman ρ**  ***P- value***  **[Confidence interval of Spearman ρ]**  **\**  **FDR-adjusted q** | **DHBS** | **Aerobactin** | **Salmochelin SX** | **RatNP-2** | **RatNP-3** | **RatNP-4** | **S100A8** | **S100A9-ac** | **S100A9-ac/m** |
| --- | --- | --- | --- | --- | --- | --- | --- | --- | --- | --- |
| **+2.28** | **DHBS** | \ | 0.4499 | 0.1808 | **0.0149** | 0.1929 | 0.1929 | **0.0033** | 0.1808 | **0.0149** |
| **+2.28** | **Aerobactin** | 0.2483  *0.3999*  [-0.4578-0.7624] | \ | 0.1808 | 0.7953 | 0.9830 | 0.9830 | 0.4144 | 0.4499 | 0.8477 |
| **+2.28** | **Salmochelin SX** | 0.9948  *0.1356*  [-0.9343-1.0000] | 0.7616  *0.1355*  [-0.5144-0.9883] | \ | 0.0861 | **0.0465** | 0.0465 | 0.1124 | 0.0769 | **0.0208** |
| **+2.28** | **RatNP-2** | **0.8051**  ***0.0044***  **[0.5236-0.9281]** | -0.0675  *0.7290*  [-0.5173-0.4116] | **0.7327**  ***0.0430***  **[0.0550-0.9482]** | \ | **0.0149** | **0.0149** | **0.0043** | **0.0033** | 0.1124 |
| **+2.28** | **RatNP-3** | 0.9763  *0.1554*  [-0.8619-1.0000] | -0.0047  *0.9830*  [-0.5264-0.5195] | **0.7677**  ***0.0194***  **[0.3022-0.9376]** | **1.0000**  ***0.0050***  **[0.9960-1.0000]** | \ | **0** | **0.0055** | 0.1124 | 0.1124 |
| **+2.28** | **RatNP-4** | 0.9763  *0.1554*  [-0.8619-1.0000] | -0.0047  *0.9830*  [-0.5264-0.5195] | **0.7677**  ***0.0194***  **[0.3022-0.9376]** | **1.0000**  ***0.0050***  **[0.9960-1.0000]** | **1.0000**  ***0***  **[1.0000-1.0000]** | \ | **0.0055** | 0.1124 | 0.1124 |
| **+2.28** | **S100A8** | **0.6977**  ***0.0003***  **[0.5764-0.7890]** | -0.1722  *0.3453*  [-0.5551-0.2709] | 0.6016  *0.0658*  [-0.0844-0.9007] | **0.9252**  ***0.0007***  **[0.8156-0.9707]** | **0.9205**  ***0.0012***  **[0.7821-0.9724]** | **0.9205**  ***0.0012***  **[0.7821-0.9724]** | \ | **0.0043** | 0.0861 |
| **+2.28** | **S100A9-ac** | 0.9810  *0.1349*  [-0.8097-1.0000] | 0.1493  *0.3947*  [-0.2803-0.5291] | **0.8086**  ***0.0342***  **[0.1568-0.9697]** | **0.9256**  ***0.0002***  **[0.8558-0.9623]** | **0.9914**  ***0.0749***  **[-0.4112-1.0000]** | **0.9914**  ***0.0749***  **[-0.4112-1.0000]** | **0.8698**  ***0.0006***  **[0.7417-0.9367]** | \ | **0.0033** |
| **+2.28** | **S100A9-ac/m** | **0.7806**  ***0.0048***  **[0.4878-0.9156]** | -0.0541  *0.8006*  [-0.5452-0.4646] | **0.6858**  ***0.0075***  **[0.4026-0.8492]** | 0.9918  *0.0720*  [-0.3738-1.0000] | 0.9929  *0.0642*  [-0.2606-1.0000] | 0.9929  *0.0642*  [-0.2606-1.0000] | **0.9993**  ***0.0410***  **[0.2577-1.0000]** | **0.9015**  ***0.0004***  **[0.8048-0.9516]** | \ |
| **-1.08** | **DHBS** | \ | 0.1177 | 0.0887 | **0.0171** | **0.0225** | **0.0171** | **0.0171** | **0.0171** | **0.0171** |
| **-1.08** | **Aerobactin** | 0.5718  *0.0817*  [-0.1294-0.8918] | \ | **0.0237** | 0.0887 | 0.0887 | 0.0887 | 0.1361 | 0.0887 | 0.0887 |
| **-1.08** | **Salmochelin SX** | 0.5657  *0.0523*  [-0.0102-0.8598] | **0.9608**  ***0.0079***  **[0.6927-0.9956]** | \ | 0.1340 | 0.1330 | 0.1330 | 0.1369 | 0.1361 | 0.1340 |
| **-1.08** | **RatNP-2** | **0.5307**  ***0.0047***  **[0.2934-0.7063]** | 0.6704  *0.0542*  [-0.0235-0.9284] | 0.6661  *0.1127*  [-0.2891-0.9567] | \ | **0.0171** | **0.0293** | 0.1330 | **0.0022** | 0.0889 |
| **-1.08** | **RatNP-3** | **0.5533**  ***0.0069***  **[0.2779-0.7447]** | 0.6986  *0.0520*  [-0.0121-0.9404] | 0.7056  *0.1029*  [-0.2725-0.9665] | **1.0000**  ***0.0040***  **[0.9974-1.0000]** | \ | **0.0293** | 0.1340 | **0.0022** | 0.0993 |
| **-1.08** | **RatNP-4** | **0.5510**  ***0.0042***  **[0.3151-0.7228]** | **0.7096**  ***0.0459***  **[0.0264-0.9410]** | 0.7135  *0.0967*  [-0.2488-0.9669] | **0.9999**  ***0.0122***  **[0.9564-1.0000]** | **0.9999**  ***0.0122***  **[0.9564-1.0000]** | \ | 0.1340 | **0.0022** | 0.0887 |
| **-1.08** | **S100A8** | **0.5181**  ***0.0039***  **[0.2986-0.6856]** | 0.5922  *0.1313*  [-0.3077-0.9329] | 0.6363  *0.1369*  [-0.3560-0.9541] | 0.9866  *0.1071*  [-0.6921-1.0000] | 0.9832  *0.1228*  [-0.7657-1.0000] | 0.9843  *0.1179*  [-0.7457-1.0000] | \ | **0.0171** | 0.1330 |
| **-1.08** | **S100A9-ac** | **0.5385**  ***0.0024***  **[0.3436-0.6889]** | 0.6573  *0.0531*  [-0.0170-0.9206] | 0.6674  *0.1323*  [-0.3629-0.9635] | **0.9498**  ***0.0001***  **[0.9021-0.9745]** | **0.9397**  ***0.0001***  **[0.8854-0.9687]** | **0.9537**  ***0.0002***  **[0.9019-0.9784]** | **0.9012**  ***0.0048***  **[0.6383-0.9758]** | \ | **0.0293** |
| **-1.08** | **S100A9-ac/m** | **0.5290**  ***0.0032***  **[0.3184-0.6899]** | 0.6793  *0.0505*  [-0.0030-0.9300] | 0.6953  *0.1211*  [-0.3411-0.9687] | 0.9941  *0.0568*  [-0.1347-1.0000] | 0.9926  *0.0662*  [-0.2904-1.0000] | 0.9945  *0.0526*  [-0.0537-1.0000] | 0.9869  *0.1049*  [-0.6783-1.0000] | **0.9999**  ***0.0122***  **[0.9564-1.0000]** | \ |

The correlations were performed at both +2.28 mm and -1.08 mm from bregma brain levels. Correlations were calculated from data obtained from the same or consecutive tissue sections using Sperman correlation analysis. Results were considered significant at *P* ≤ 0.05 (values in red italic). The strength and direction of the association between variables are expressed as Sperman ρ-values (values in bold). To account for multiple comparisons, *P* values were adjusted using the Benjamini–Hochberg false discovery rate (FDR) procedure; FDR-adjusted q values are reported, and correlations with FDR q ≤ 0.05 (values in bold) were considered statistically significant after correction. DHBS, dihydroxybenzoylserine; RatNP, rat neutrophil peptide; ac, acetylation; m, methylation.

**Table S5. Regional ranking of host–siderophore dissociation in infected brain tissue.**

| **Brain level from bregma (mm)** | **Brain region** | **Ranking of the host-siderophore dissociation** | **Mean of host score ± SD** | **Mean of siderophore score ± SD** | **Mean of host-siderophore score ± SD** |
| --- | --- | --- | --- | --- | --- |
| +2.28 | Artery | 1 | 1.7252 ± 0.7174 | 0.0142 ± 0.2588 | 1.7110 ± 0.5537 |
| +2.28 | LS | 2 | -0.0883 ± 0.3879 | -0.0833 ± 0.2515 | -0.0050 ± 0.2639 |
| +2.28 | Ctx | 3 | -0.6107 ± 0.0876 | -0.4109 ± 0.0051 | -0.1998 ± 0.0836 |
| +2.28 | CPu | 4 | -0.6611 ± 0.0387 | -0.3747 ± 0.0436 | -0.2864 ± 0.0508 |
| +2.28 | AcbN | 5 | -0.6474 ± 0.0602 | -0.3501 ± 0.0485 | -0.2973 ± 0.0575 |
| +2.28 | cc | 6 | -0.6159 ± 0.0514 | -0.2539 ± 0.1305 | -0.3620 ± 0.1036 |
| +2.28 | LV | 7 | 0.7751 ± 0.9151 | 1.4080 ± 2.1893 | -0.6329 ± 1.6355 |
| -1.08 | D3V | 1 | 1.5560 ± 1.3319 | 0.4816 ± 0.7754 | 1.0744 ± 1.1882 |
| -1.08 | 3V | 2 | 1.6371 ± 0.7524 | 1.0850 ± 1.0087 | 0.5521 ± 1.5352 |
| -1.08 | ATh | 3 | -0.4282 ± 0.1499 | -0.4549 ± 0.1034 | 0.0267 ± 0.1496 |
| -1.08 | fi+vhc | 4 | -0.3533 ± 0.2105 | -0.3509 ± 0.2768 | -0.0023 ± 0.3376 |
| -1.08 | Ctx | 5 | -0.5694 ± 0.0565 | -0.5412 ± 0.0122 | -0.0283 ± 0.0591 |
| -1.08 | AHth | 6 | -0.4065 ± 0.1105 | -0.3327 ± 0.1033 | -0.0738 ± 0.1075 |
| -1.08 | CPu | 7 | -0.6103 ± 0.0147 | -0.5192 ± 0.0179 | -0.0911 ± 0.0232 |
| -1.08 | GP | 8 | -0.6245 ± 0.0158 | -0.4784 ± 0.0514 | -0.1460 ± 0.0501 |
| -1.08 | cc | 9 | -0.5174 ± 0.0934 | -0.3356 ± 0.2883 | -0.1818 ± 0.2887 |
| -1.08 | LV | 10 | 0.5649 ± 0.6316 | 1.5476 ± 1.7188 | -0.9827 ± 1.5169 |

Ranking of brain regions according to the difference between composite host-response and composite siderophore scores in infected animals *(n* = 5). Analyses were performed separately for the +2.28 mm and −1.08 mm bregma levels. For each analyte, regional MSI abundances were standardized using z-score normalization within each bregma level. The mean host score was calculated as the average z-score of RatNP-2, RatNP-3, RatNP-4, S100A8, acetylated S100A9 and acetylated/methylated S100A9. The mean siderophore score was calculated as the average z-score of 2,3-dihydroxybenzoylserine, aerobactin and salmochelin SX. The host–siderophore score was calculated as the difference between the mean host score and the mean siderophore score. Regions were ranked according to the magnitude of the host–siderophore score within each bregma level. Positive values indicate relative enrichment of host antimicrobial markers, whereas negative values indicate relative enrichment of siderophore signals. SD, standard deviation; artery, cerebral artery; LS, lateral septum; Ctx, cortex; CPu, caudate putamen; AcbN, accumbens nucleus; cc, corpus callosum; LV, lateral ventricle; D3V, dorsal third ventricle; 3V, third ventricle; ATh, anterior thalamus; AHth, anterior hypothalamus; fi+vhc, fimbria of hippocampus; GP, globus pallidus.

**Table S6.** **List of assigned endogenous peptides with high mass accuracy as determined by MALDI FTICR-MSI experiments.**

| **Species ID** |  |  |  | **Molecular**  **Formula** | **Molecular ion** | **Measured *m/z*** | **Theor. *m/z*** | **Error ppm** | **Multiple comparison ctrl vs. inf**  ***P*-value at +2.28/-1.08 mm level** | **Fold change ctrl vs. inf at the +2.28/-1.08 mm level** | **UniProt ID** |
| --- | --- | --- | --- | --- | --- | --- | --- | --- | --- | --- | --- |
| **Peptide precursor** | **Peptide name** | **Peptide sequence** | **Postrans. modific.; -S-S- bond** |  |  |  |  |  |  |  |  |
| Proenkephalin (PEnk) | Met‐Enk‐Arg‐Phe (263-269) | YGGFMRF |  | C42H56N10O9S | [M+H]^+^ | 877.4027 | 877.4025 | 0.20 | 0.3910/0.1185 | n.a./ 4.3 | P04094 |
|  | Met‐Enk‐Arg‐Gly-Leu (188-195) | YGGFMRGL |  | C41H61N11O10S | [M+H]^+^ | 900.4407 | 900.4396 | 1.14 | n.a./ 0.0817 | n.a./ 4.7 | P04094 |
|  | PEnk (220-229) | GRPEWWMDYQ |  | C63H82N16O17S | [M+H]^+^ | 1367.5836 | 1367.5837 | -0.07 | 0.4732/0.3923 | 4.2/ 1.7 | P04094 |
|  | PEnk (198‐209) | SPQLEDEAKELQ |  | C58H95N15O24 | [M+H]^+^ | 1386.6795 | 1386.6753 | 2.77 | 0.2191/0.0815 | 6.3/ 2.3 | P04094 |
|  | PEnk (219-229) | VGRPEWWMDYQ |  | C68H91N17O18S | [M+H]^+^ | 1466.6570 | 1466.6521 | 3.34 | 0.1633/**0.0067** | n.a./ 1.9 | P04094 |
|  | PEnk (114-133) | MDELYPVEPEEEANGGEILA |  | C95H145N21O37S | [M+H]^+^ | 2204.9933 | 2204.9904 | 1.35 | 0.4999/0.1263 | 1.6/ 2.1 | P04094 |
|  | PEnk (212-229) | YGGFMRRVGRPEWWMDYQ |  | C107H148N30O26S2 | [M+H]^+^ | 2334.0759 | 2334.0695 | 2.74 | **0.0076/**0.5177 | 7.6/0.8 | P04094 |
|  | PEnk (239-260) | FAESLPSDEEGESYSKEVPEME |  | C106H157N23O44S | [M+H]^+^ | 2489.0530 | 2489.0548 | -0.73 | 0.1929/0.1199 | 3.4/2.5 | P04094 |
|  |  |  |  |  | [M+K]^+^ | 2527.0108 | 2527.0107 | 0.03 | 0.5470/0.0996 | 1.4/1.6 |  |
|  | PEnk (239-260) | FAESLPSDEEGESYSKEVPEME | Phosph. (1x) | C106H158N23O47SP | [M+H]^+^ | 2569.0285 | 2569.0212 | 2.87 | 0.1354/0.1272 | 2.9/1.9 | P04094 |
|  |  |  |  |  | [M+K]^+^ | 2606.9858 | 2606.9770 | 3.4 | 0.1045/0.0955 | 5.5/1.9 |  |
|  | PEnk (143-185) | DADEGDTLANSSDLLKELLGTGDNRAKDSHQQESTNNDEDSTS |  | C181H290N56O84 | [M+H]^+^ | 4593.0173 | 4593.0215 | -0.92 | 0.6708/**0.0177** | 1.2/1.9 | P04094 |
| Cerebellin 1  Precursor protein | Cerebellin (1-15) | SGSAKVAFSAIRSTN |  | C63H106N20O22 | [M+H]^+^ | 1495.7939 | 1495.7863 | 5.08 | n.a./ 0.1896 | n.a./ 0.5 | P63182 |
|  | Cerebellin 1 (57-72) | SGSAKVAFSAIRSTNH |  | C69H113N23O23 | [M+H]^+^ | 1632.8465 | 1632.8452 | 0.76 | 0.2135/0.1981 | 2.1/0.7 | P63182 |
| ProSAAS | Big LEN (245-260) | LENSSPQAPARRLLPP |  | C76H128N24O23 | [M+H]^+^ | 1745.9662 | 1745.9656 | 0.33 | **0.0016/0.0253** | 8.7/1.7 | Q9QXU9 |
|  |  |  |  |  | [M+DHB-H2O+Na]+ | 1903.9664 | 1903.9637 | 1.44 | **0.0016/**0.8926 | 0.5/1.0 |  |
|  | Little SAAS (42-49) | SLSAASAPLAETSTPLRL |  | C77H133N21O27 | [M+H]^+^ | 1784.9744 | 1784.9752 | -0.48 | 0.0672/**0.0124** | 0.04/1.54 | Q9QXU9 |
|  |  |  |  |  | [M+K]^+^ | 1822.9316 | 1822.9311 | 0.25 | 0.3739/**0.0097** | n.a./ 1.9 |  |
|  | PEN-20 (221-240) | AVDQDLGPEVPPENVLGALL |  | C91H148N22O31 | [M-H2O+H]+ | 2028.0748 | 2028.0648 | 4.97 | **0.0051/**0.7204 | 0.02/1.6 | Q9QXU9 |
| Thymosin beta-4 | Thymosin beta-4 (2-44) | SDKPDMAEIEKFDKSKLKKTETQEKNPLPSKETIEQEKQAGES | Acetyl. (1x) | C212H350N56O78S | [M+H]^+^ | 4961.4917 | 4961.4936 | -0.39 | 0.6594/0.5820 | 1.0/1.0 | P62329 |
|  |  |  |  |  | [M+Na]^+^ | 4983.4840 | 4983.4755 | 1.69 | 0.7913/**0.0216** | 1.00.8 |  |
|  |  |  | Phosph. (1x) | C210H349N56O80SP | [M+H]^+^ | 4999.4566 | 4999.4494 | 1.46 | 0.5317/0.7182 | 1.0/1.0 | P62329 |
|  |  |  | Phosph. (1x), Acetyl. (1x) | C217H360N57O82S2P | [M-H2O+H]+ | 5154.4908 | 5154.4898 | 0.18 | 0.0746/0.5729 | 1.3/1.0 | P62329 |
|  |  |  | Phosph. (2x), Acetyl. (2x) | C213H357N57O83SP2 | [M+H]^+^ | 5136.4843 | 5136.4735 | 2.11 | **0.0031/**0.7732 | 0.7/1.0 | P62329 |
| Thymosin beta-10 | Thymosin beta-10 (2-44) | ADKPDMGEIASFDKAKLKKTETQEKNTLPTKETIEQEKRSEIS | Phosph. (1x), Acetyl. (3x) | C215H358N57O81SP | [M+K]^+^ | 5098.5047 | 5098.5177 | -2.57 | **0.0209/**0.9373 | 0.7/1.0 | P63312 |
| Neutrophil antibiotic peptide NP-3A/B | RatNP-3A/B (59-87) | CSCRTSSCRFGERLSGACRLNGRIYRLCC | -S-S- (3x) | C130H214N48O39S6 | [M+H]^+^ | 3264.4660 | 3264.4634 | 0.45 | **0.0055/0.0103** | 0.004/0.0008 | Q62713/Q9Z1F1 |
|  |  |  |  |  | [M+Na]^+^ | 3286.4404 | 3286.4454 | -1.51 | **0.0035/0.0142** | 0.001/0.00009 |  |
|  |  |  |  |  | [M+K]^+^ |  | 3302.4845 |  | **0.0067/0.0089** | 0.002/0.0002 |  |
| Defensin alpha 4 | RatNP-4 (63-93) | ACYCRIGACVSGERLTGACGLNGRIYRLCCR | -S-S- (3x) | C136H223N47O39S6 | [M+H]^+^ | 3331.5373 | 3331.5308 | 1.95 | **0.0060/0.0087** | 0.004/0.0008 | Q62714 |
|  |  |  |  |  | [M+Na]^+^ |  | 3353.5128 |  | **0.0031/0.015** | 0.002/0.001 |  |
|  |  |  |  |  | [M+K]^+^ | 3369.4920 | 3369.4867 | 1.57 | **0.0068/0.0074** | 0.003/0.0001 |  |
| Neutrophil antibiotic peptide NP-2 | RatNP-2 (63-94) | VTCYCRSTRCGFRERLSGACGYRGRIYRLCCR | -S-S- (3x) | C155H248N56O42S6 | [M+H]^+^ | 3758.7451 | 3758.7389 | 1.67 | **0.0084/0.0091** | 0.005/0.003 | Q62715 |
|  |  |  |  |  | [M+K]^+^ |  | 3796.6948 |  | **0.0086/0.0063** | 0.002/0.0003 |  |

Peptides detected in control (ctrl, *N* = 4) and infected (inf, *N* = 5) brain tissue sections were analyzed at +2.28 mm and −1.08 mm relative to bregma. Peptides were detected in positive-ion mode predominantly as protonated species, with a limited number of potassiated and sodiated ions. Peptide abundances were normalized using root mean square (RMS) normalization, except for the antimicrobial peptides RatNP-2, RatNP-3, and RatNP-4, which were normalized to the peak area of mouse hepcidin-25. Relative peptide abundances between control and infected tissue sections were compared separately at bregma level using an unpaired t-test. Differences were considered statistically significant at P ≤ 0.05 (indicated in bold). Fold changes were calculated as the ratio of the mean peptide abundance in infected versus control tissue sections.

**Table S7. Brain regional quantitation of rat antimicrobial peptides (RatNP1-3), aerobactin, 2,3-dihydroxybenzoylserine and salmochelin SX in control (*n* = 4) and infected (*n* = 5) brain tissue sections.**

| **Brain regions** | **Level from bregma [mm]** | **RatNP-2 [µg/g]** | | **RatNP-3 [µg/g]** | | **RatNP-4 [µg/g]** | | **Aerobactin** | | **DHBS** | | **Salmochelin SX** | |
| --- | --- | --- | --- | --- | --- | --- | --- | --- | --- | --- | --- | --- | --- |
|  |  | **Ctrl** | **Inf** | **Ctrl** | **Inf** | **Ctrl** | **Inf** | **Ctrl** | **Inf** | **Ctrl** | **Inf** | **Ctrl** | **Inf** |
| **AcbN** | +2.28 | nd | det | nd | det | nd | det | nd | det | nd | nd | nd | nd |
| **artery** | +2.28 | nd | **15.9** | nd | **30.6** | nd | **29.5** | nd | det | nd | nd | nd | nd |
| **cc** | +2.28 | nd | det | nd | det | nd | det | nd | det | nd | nd | nd | nd |
| **CPu** | +2.28 | nd | det | nd | det | nd | det | nd | det | nd | nd | nd | nd |
| **Ctx** | +2.28 | nd | det | nd | det | nd | det | nd | det | nd | nd | nd | nd |
| **LS** | +2.28 | nd | **5.8** | nd | **9.9** | nd | **9.5** | nd | det | nd | nd | nd | nd |
| **LV** | +2.28 | nd | **9.3** | nd | **15.8** | nd | **15.6** | nd | det | nd | nd | nd | nd |
| **3V** | -1.08 | nd | **12.3** | nd | **23.8** | nd | **21.9** | nd | det | nd | det | nd | nd |
| **AHth** | -1.08 | nd | det | nd | det | nd | det | nd | det | nd | det | nd | nd |
| **ATh** | -1.08 | nd | det | nd | **5.3** | nd | det | nd | det | nd | det | nd | nd |
| **cc** | -1.08 | nd | det | nd | det | nd | det | nd | det | nd | det | nd | nd |
| **CPu** | -1.08 | nd | det | nd | det | nd | det | nd | det | nd | det | nd | nd |
| **Ctx** | -1.08 | nd | det | nd | det | nd | det | nd | det | nd | det | nd | nd |
| **D3V** | -1.08 | nd | **13.0** | nd | **28.3** | nd | **26.2** | nd | det | nd | det | nd | nd |
| **fi+vhc** | -1.08 | nd | det | nd | **6.8** | nd | **6.2** | nd | det | nd | det | nd | nd |
| **GP** | -1.08 | nd | det | nd | det | nd | det | nd | det | nd | det | nd | nd |
| **LV** | -1.08 | nd | **9.3** | nd | **18.3** | nd | **17.0** | nd | det | nd | det | nd | nd |

The quantitation of RatNPs and siderophores was performed using MALDI-qMSI approach. The method validation parameters are shown in Fig. S2 and Table S2. AcbN, accumbens nucleus; artery, cerebral artery; cc, corpus callosum; CPu, caudate putamen; Ctx, cortex; LS, lateral septum; LV, lateral ventricle; 3V, third ventricle; AHth, anterior hypothalamus; ATh, anterior thalamus; D3V, dorsal third ventricle; fi+vhc, fimbria of hippocampus; GP, globus pallidus; Ctrl, control; Inf, infected; nd, not detected (value below calculated limit of detection); det, detected (value between limit of detection and limit of quantitation).

**Table S8. Sequences and final concentrations of qPCR primers and hydrolysis probe used for qPCR.**

| **Primer/probe** | **Sequence** | **Concetration (nM)** |
| --- | --- | --- |
| Forward primer *uid*F | 5’-GCGGCAAAGGATTCGATAAC-3’ | 300 |
| Reverse primer *uid*R | 5’-TCTCTTCAGCGTAAGGGTAATG-3’ | 300 |
| Hydrolysis probe *uid*P | 5’-FAM-TCACGCACTAATGGACTGGATTGGG-TAMRA-3’ | 250 |
